# Intraspecies associations from strain-rich metagenome samples

**DOI:** 10.1101/2025.02.07.636498

**Authors:** Evan B. Qu, Jacob S. Baker, Laura Markey, Veda Khadka, Chris Mancuso, Delphine Tripp, Tami D. Lieberman

## Abstract

Genetically distinct strains of a species can vary widely in phenotype, reducing the utility of species-resolved microbiome measurements for detecting associations with health or disease. While metagenomics theoretically provides information on all strains in a sample, current strain-resolved analysis methods face a tradeoff: *de novo* genotyping approaches can detect novel strains but struggle when applied to strain-rich or low-coverage samples, while reference database methods work robustly across sample types but are insensitive to novel diversity. We present PHLAME, a method that bridges this divide by combining the advantages of reference-based approaches with novelty awareness. PHLAME explicitly defines clades at multiple phylogenetic levels and introduces a probabilistic, mutation-based, framework to accurately quantify novelty from the nearest reference. By applying PHLAME to publicly available human skin and vaginal metagenomes, we uncover previously undetected clade associations with coexisting species, geography, and host age. The ability to characterize intraspecies associations and dynamics in previously inaccessible environments will propel new mechanistic insights from accumulating metagenomic data.

## Introduction

Assessing whether microorganisms are statistically associated with sample features, such as disease state, environmental conditions, or species richness, is a fundamental line of inquiry in microbiome science. These relationships have been explored extensively at the species level and above, revealing generalizable associations of bacterial taxa with host disease^1,2^, diet^3^, life stage^4,5^, and more. However, given that strains of the same bacterial species can possess genetically-encoded phenotypic differences, there is increasing interest^6–9^ in identifying associations between intraspecies population structure and sample features of microbiomes.

The rapid accumulation of publicly available metagenomic data has the potential to enable new, well-powered genetic association studies within microbial species. However, available strain-resolved metagenomic methods face significant limitations when samples contain high strain richness or have low coverage. The most popular approaches use metagenome-assembled genomes (MAGs)^10^ or alignment-based^11,12^ methods to reconstruct a single dominant genotype per species, per sample and are thus limited in samples with high intraspecies diversity^11–13^. A second class of methods leverages covariation between genomic elements in related samples to infer multiple *de novo* genotypes, but requires high sequencing depth per strain and several related samples for each profiled community^14,15^. As such, the discovery of strain-level associations has been mostly restricted to environments where intraspecies diversity is low, such as the human gut microbiome^8,16^, and there are fewer known strain-feature relationships in environments with high strain richness and low biomass, including human skin and the female genital tract.

Methods that query metagenomes against a collection of curated reference genomes (reference database methods) have shown strong performance at low sequencing depth and in the presence of many closely-related strains^7–9^. The rate of high-quality reference genomes available for such approaches continues to rise, with individual studies now reporting thousands of bacterial isolates cultured from the human gut^17^ or skin^18,19^. However, these approaches are still limited to characterizing diversity represented in the reference set, which may be missing portions of intraspecies diversity due to collection or culturing biases. For analyses at very fine evolutionary resolutions, it is expected that new samples will always have strain diversity not represented in the reference set, as bacteria constantly accumulate genetic changes over time^20^. Current reference database methods do not account for this uncertainty and often report novel strains as combinations of related genomes present in the reference database (Fig. 1A).

**Figure 1:**
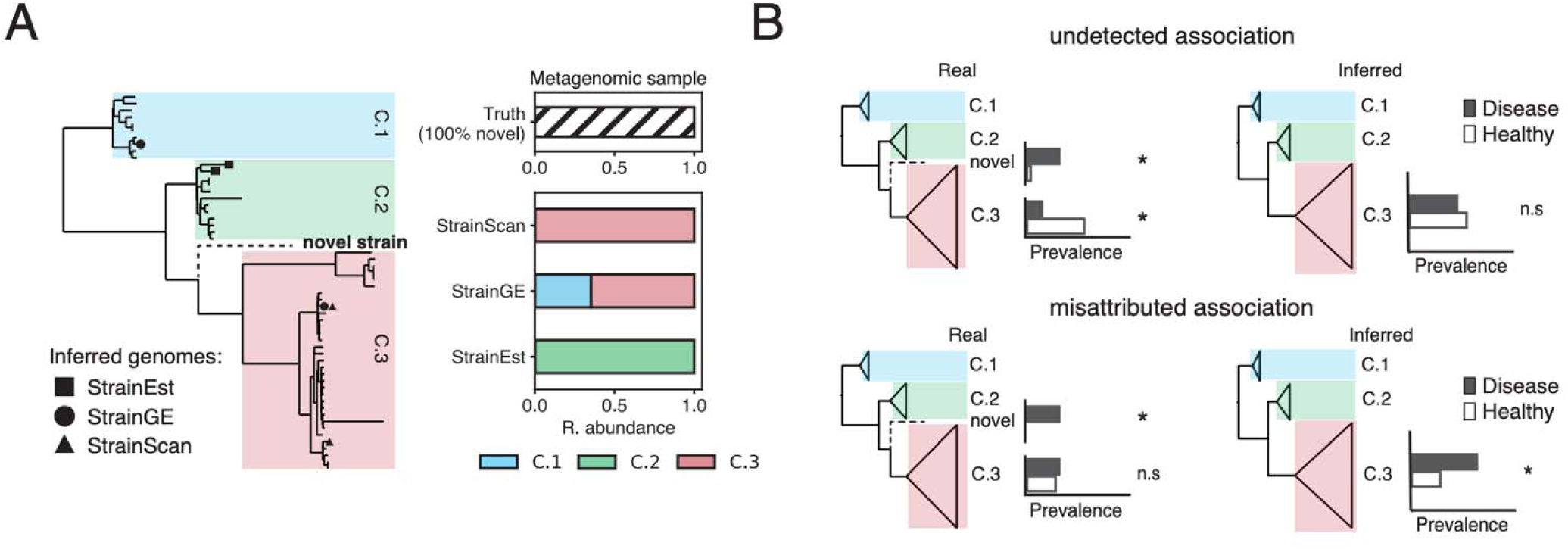
Reference-based methods for strain profiling represent novel diversity as combinations of known genomes. **(A)** Left: Phylogenetic representation of a section of an intraspecies reference database, labeled with major within-species clades. We examined the output of 3 different methods when given an input metagenome only containing only a “novel” strain (held out from the database and shown as the dashed branch). Each dot represents a genome output by its associated method. Right: Taxonomic bar plots showing the clade-level composition reported by different methods. Representing novel diversity as known genomes can lead to inconsistent clade-level results between methods. **(B)** Two hypothetical scenarios where representing novel diversity as known strains might convolute the detection of intraspecies associations. For each clade, the prevalence of the clade in a diseased and healthy cohort is shown; stars indicate hypothetically significant (left) or measured significant (right) associations. In the top row, a novel clade and a known clade have inverse associations (e.g., one is associated with disease, and the other is associated with health). If both clades are detected as the known clade, both associations are lost. In the bottom row, an association of the novel clade with disease is detected as an association driven by a known clade, which is potentially misleading if the novel and known clade have different phenotypic characteristics.

Reporting novel diversity as known reference genomes can obscure or misattribute real taxonomic associations (Fig 1B). For example, given a novel clade and a known clade with inverse associations (e.g., one is associated with disease and the other is associated with health), both associations will be masked if the novel clade is reported as a member of the known clade. Alternatively, an association driven by a novel clade can be misattributed as an association driven by a known clade, which is misleading if the two clades have different phenotypic characteristics.

Here, we present PHLAME – a novelty-aware approach to classify intraspecies variation and detect associations using reference databases. PHLAME defines multiple hierarchical groupings of clades from the species phylogeny and implements a Bayesian zero-inflated negative binomial model to both infer clade abundances and quantify support for either a known or novel clade in a sample. By identifying highly conserved core genome mutations that accumulate at a clock-like rates^21^, PHLAME accurately quantifies the phylogenetic novelty of strains in metagenome samples. Using both simulations and real data, we highlight new analyses that PHLAME enables, including the ability to identify samples with abundant novel strain diversity for culturomics efforts. Finally, we demonstrate the utility of PHLAME in detecting associations in strain-rich human skin and vaginal microbiome samples.

## Results

### Reference-based methods for strain profiling present novel diversity as combinations of known genomes

The behavior of different strain-level reference database methods in the presence of novel diversity has not been compared in the literature. We used simulations to examine how novel diversity affects the behavior of three published methods that characterize strain mixtures in metagenomic samples (StrainEst^22^, StrainGE^23^, and StrainScan^24^). Performance in the presence of novel diversity is typically evaluated by excluding single genomes from reference databases^22–24^, but existing works have not examined scenarios where entire phylogenetic clades are missing, as would be expected from biased or incomplete cultivation efforts. To test the impact of systematically missing phylogenetic diversity on classification, we generated phylogenies for three species: *C. acnes*, *S. epidermidis*, and *E. coli* and defined well-supported intraspecies clades within each species (Methods). We iteratively held out a single clade and all its descendants from the reference database and compared the performa ce of each method when faced with a sample comprised of a single genome from the held-out clade (Methods).

Although the true genome was never present in the reference database, methods reported one or more detected genomes in most instances (50%, 100%, and 77% of all simulations for StrainEst, StrainGE, and StrainScan, respectively). Held-out genomes that were more novel (that is, more genetically distant to the remaining reference set) were less likely to be classified by StrainEst, but not by StrainScan or StrainGE (Fig. S1C). When strains were reported, the most genetically similar reference was not returned for the majority of cases (> 90% of simulations for all methods). To quantify assignment error more precisely, we calculated Excess Distance, which compares the distance between the true held-out genome with the output genome(s) to the distance between the true held-out genome and its closest reference in the database (Methods). There was no significant difference in Excess Distance between different methods (Fig. S1A). The median Excess Distance across the core genome for each species was 171 SNPs for *C. acnes*, 1,068 SNPs for *S. epidermidis*, and 1,724 SNPs for *E. coli*.

The genomes output by different methods on the same simulation were often located in different parts of the phylogeny. We quantified the phylogenetic consistency of methods with one another by calculating the mean pairwise UniFrac distance (mp-UniFrac) of each method’s output (Fig. S1B). For *E. coli*, this metric was only calculated between StrainGE and StrainScan, as StrainEst did not return an output for a majority of simulations. The median mp-UniFrac between method predictions was highest for *E. coli* (0.32), followed by *S. epidermidis* (0.28) and *C. acnes* (0.03), correlating with both median Excess Distance and the overall genetic diversity of each species. Together, these simulations highlight existing challenges of accurate strain-level classification when reference databases are incomplete.

### PHLAME quantifies the divergence of novel strains in metagenomes

We present PHLAME, a reference database method with two critical features: (1) classification at defined hierarchical evolutionary groupings called clades (Methods); and (2) a probabilistic model that explicitly quantifies phylogenetic novelty for multiple strains in a metagenomic sample. PHLAME uses an alignment and single-nucleotide variant (SNV) based approach to identify alleles that are specific to and unanimous within all reference genomes within a clade (clade-specific alleles). This clade-based approach operates at the natural level for detecting intraspecies associations while increasing sensitivity by including markers common to multiple strains. Because it is often not obvious what phylogenetic resolution to detect associations at, PHLAME reports a frequency for all well-supported clades in a sample simultaneously.

PHLAME leverages the fact that novel strains share an underlying evolutionary history with known clades of the same species, and thus will share some, but not all, clade-specific alleles with its closest relative (Fig. 2A). We refer to this degree of shared history between a novel strain and its closest clade in the database as Divergence (DV_b_), which is calculated with respect to a specific branch on the phylogeny (*b*). Low DV_b_ values support the existence of a known clade in a reference database, while high DV_b_ values support the existence of a related but novel clade.

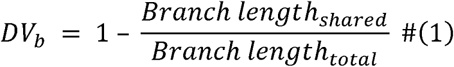

**Figure 2:**
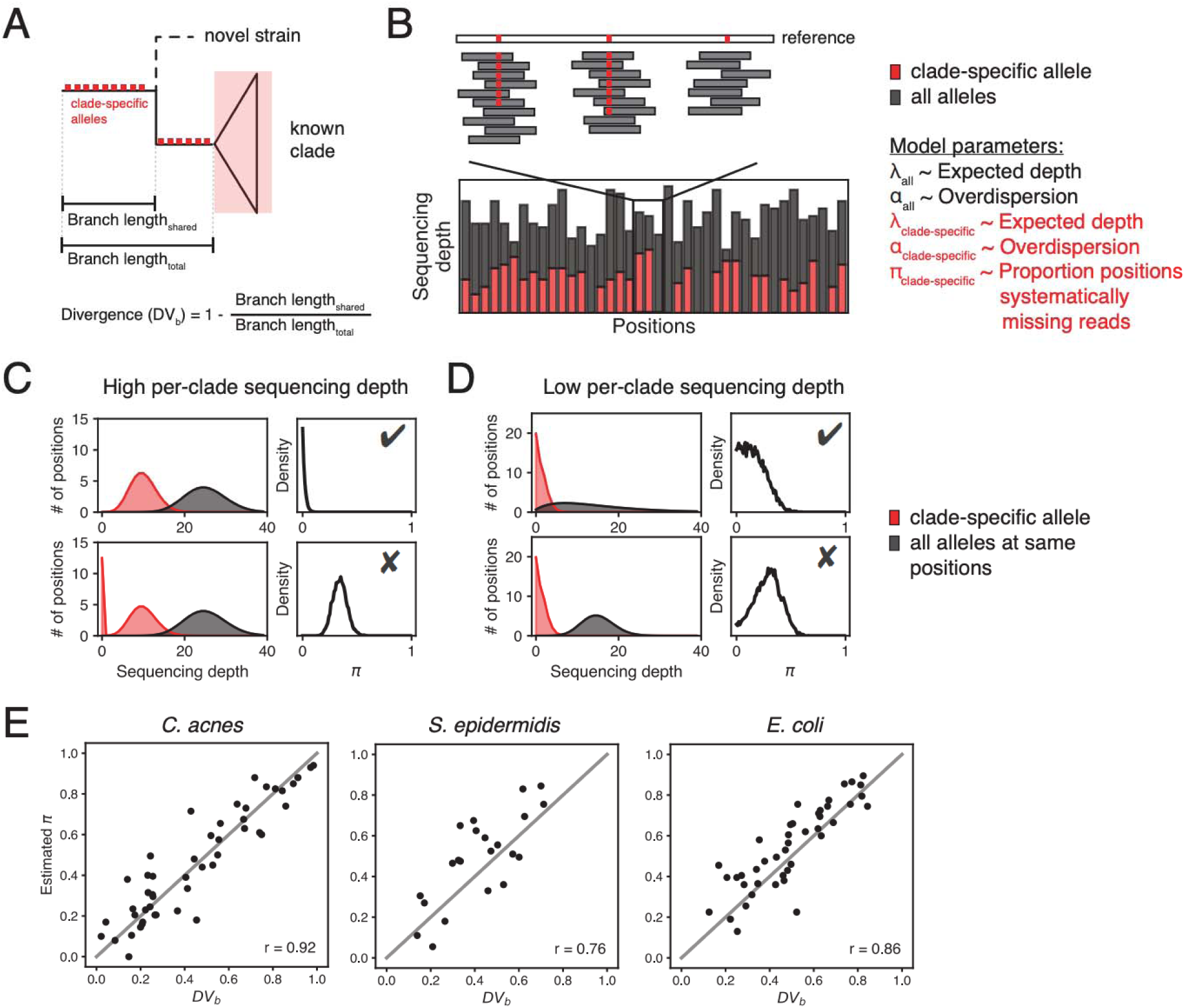
PHLAME quantifies the divergence of novel strains in metagenomes. **(A)** Diagram showing a novel strain and its closest relative in a reference database. PHLAME assumes that a novel strain will share clade-specific mutations (red boxes) until it diverges with its closest relative. Divergence (DV_b_) is a metric that measures the degree of relatedness between a novel strain and its closest relative in a database. When a clade is truly present, a sample should have all the mutations that occurred on the branch leading up that clade (clade-specific alleles). Novel strain(s) should contain only the subset of clade-specific alleles that occurred before its divergence point. **(B)** A view of allele counts across informative positions (positions containing a clade-specific mutation) for a given clade. Counts supporting the clade-specific allele are shown in red; counts supporting all other alleles are shown in grey. Discontinuities in the number of counts supporting clade-specific alleles indicate when a sample is systematically missing clade-specific mutations. **(C)** PHLAME uses a Bayesian zero-inflated model to infer the proportion of positions that systematically have zero reads from the distribution of read depths. Estimates on the proportion of positions that systematically have zero reads are represented as a posterior distribution over the parameter π. At high per-clade sequencing depths, discontinuities in read depth are obvious and it is easy to distinguish samples supporting a known clade (top) from those supporting a novel clade (bottom). **(D)** At low per-clade sequencing depths, prior information of the dispersion of the reads (gray) is critical for distinguishing between scenarios. Thresholds over the π posterior density can be used to differentiate between known (⍰) and novel () clades. **(E)** Maximum a posteriori estimates of π obtained from classifying a single held-out genome (subsampled to 10X coverage across the reference) correlate well with true DV_b_ values as determined from the full tree (Pearson correlation coefficients shown within each plot).

The key signal for DV_b_ in metagenomic samples comes from evidence of missing clade-specific markers. Many reference database methods use a heuristic centered around missing markers to reject detections of novel genomes – for example, by requiring the proportion of missing markers in a sample to be below a threshold^23^. However, marker absence counts are not a true measure of DV_b_ because markers can be missing by chance due to either low sequencing depth or high read count variance (dispersion) (Fig. S3A, S3B). While systematically missing alleles are easily identified at high per-species sequencing depths (Fig. 2C), at low sequencing depths it is difficult to determine whether zero counts are generated from low coverage, overdispersion, or high values of DV_b_ (Figs. 2D, S3C). Simple parameter estimation approaches, including both marker absence counts and maximum likelihood zero-inflation models, yield inaccurate estimates of DV_b_ under these conditions (Fig. S3D, E).

We implement a two-step model that leverages the allelic diversity across a set of positions to reduce uncertainty when estimating DV_b_ (Methods). The first step models the read depth across all alleles at informative positions as a negative binomial distribution with two parameters: expected depth (λ_all_) and excess dispersion (α_all_). The second step models read depth across only clade-specific alleles as a zero-inflated negative binomial distribution with three parameters: expected depth (λ_clade-specific_), excess dispersion (α_clade-specific_), and the proportion of systematically zero counts (π_clade-specific_, or just π). Critically, the posterior estimate over α_all_ from the first step is used to inform a prior over α_clade-specific_ and constrain possible values of π_clade-specific_and λ_clade-specific_(Fig. 2D). We demonstrate using counts simulations that incorporating prior information on the dispersion of reads reduces error in DV_b_ estimation, regardless of the inference approach used (Fig. S3D). We implement a Bayesian sampling algorithm to recover full posterior probabilities over each parameter (Supplemental Methods), as well as a maximum likelihood approach that offers faster runtime with minimal performance loss (Fig. S5, Fig. S10). This strategy thereby reduces uncertainty by independently estimating dispersion (from reads with any allele) and divergence (from reads with clade-specific alleles) at the same genomic loci.

To validate our conceptual model and PHLAME’s ability to approximate DV_b_, we performed simulations in which we iteratively held out clades in a species phylogeny from the PHLAME reference database. We used each held-out database to profile single held-out genomes subsampled to different coverages (Methods). At 10X coverage, the average Pearson correlation coefficient between metagenomic estimates of π and true values of DV_b_ was 0.85 (Fig. 2E), although significant correlations were recovered at per-clade coverages as low as 0.5X (Fig. S4). The primary detection threshold in PHLAME is based on DV_b_ and can be adjusted to reflect study-specific beliefs on how related a genome must be to be considered a member of a clade. At very low sequencing depths, the posterior distribution of π captures uncertainty in estimate of DV_b_ and can be used to quantify the degree of support for either a known or novel clade.

In addition, PHLAME avoids errors in relative abundance estimation introduced by novel diversity. The zero-inflated model better estimates clade abundances by ensuring that systematically missing markers are not average into the inferred relative abundance. Moreover, PHLAME does not force clade frequencies to sum to one, as such normalization will distort relative abundance estimates if novel strains make up a substantial fraction of the sample. PHLAME thereby provides a quantitative measure of both known and novel intraspecies diversity within a sample, with the unclassified relative abundance at a given phylogenetic resolution representing the estimated proportion of a sample comprised of novel diversity.

### PHLAME achieves improved performance in multiple-strain metagenomes

We benchmarked PHLAME’s ability to identify clades in synthetic metagenomes against comparable methods. We assessed performance for two species, *C. acnes* and *S. epidermidis,* using both a perfect reference database, where all genomes that could be in a sample are included in the database (Fig. 3A), as well as a database where 25% of the clades were randomly held out (Fig. 3B). We created complex synthetic metagenomes by combining reads from five sequenced genomes included in the database at random abundances, along with genomes from nine other species commonly found in the human skin microbiome (Supplemental Table 1). We varied total coverage of the focal species from 0.1X to 20X across the reference genome while keeping the rest of the background metagenome the same. When clades were held out from the reference database, synthetic metagenomes also contained reads from 5 randomly chosen held-out genomes for an additional 25% of the total reads of the 5 original genomes.

**Figure 3:**
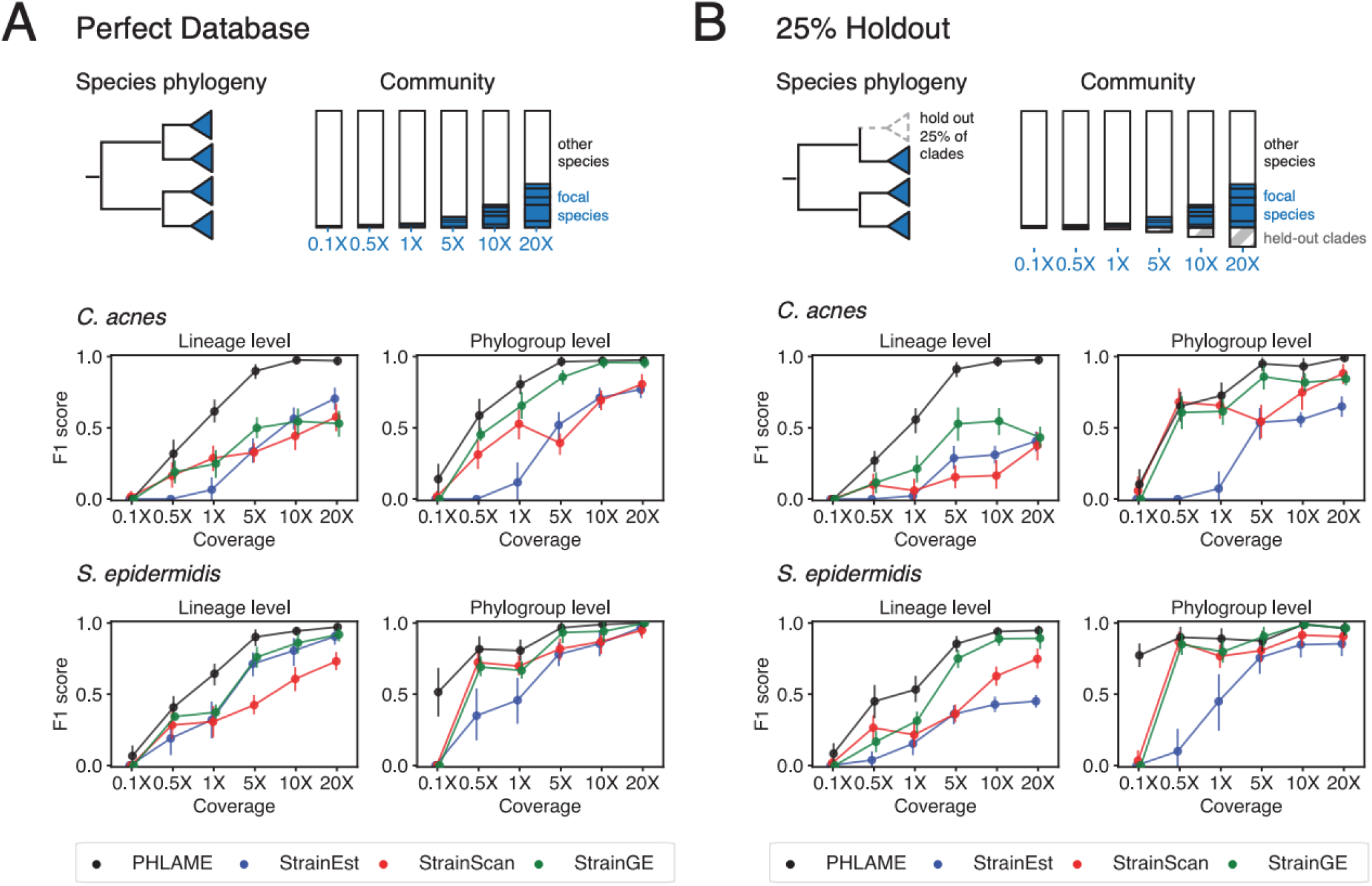
PHLAME achieves improved performance in complex multiple-strain metagenomes. In each synthetic metagenome, five random strains of the species under comparison were combined at defined abundances, along with genomes of other species. Coverage of the species under comparison varied up to a maximum of 20X, corresponding to a maximum of 20% of the community. (A) F1 score given a perfect database, where all genomes in a sample are also found in the reference database. Dots and bars represent mean and 95% confidence interval of F1 score across 15 simulations. (B) F1 score given a database where 25% of clades are randomly held out from the reference database. In each synthetic metagenome, 5 known strains were chosen from the clades included in the reference database, as well as five random held-out genomes comprising 25% of the total coverage of the five known strains. Recall and precision were determined using only the known genomes.

We evaluated performance using F1 score (Fig. 3) and L2 distance (Fig. S7) at two separate phylogenetic resolutions: coarse-grained phylogroups and fine-grained lineages. Phylogroups were defined according to established typing schemes for each species^25,26^ (Fig. S2); lineages represent finely-resolved clades separated by fewer than 100 mutations across the core genome^18,19^. For StrainEst, StrainGE, and StrainScan, we tested both a complete database and a de-replicated database containing a single genome per lineage (Fig. S5); we report results from the better performing database for each set of simulations in (Fig. 3).

Overall, PHLAME achieved improved precision, recall, and L2 distance compared to other methods (Fig. 3, Fig. S6), and these results were consistent across reasonable thresholds for detection (Fig. S9). Notably, PHLAME achieved near-perfect precision across simulations (average 0.99 across all simulations and coverages, compared to 0.63 for StrainEst, 0.84 for StrainGE, 0.80 for StrainScan). When depth of the focal species was at least 5X, PHLAME also had high recall (0.94 across all simulations). The largest improvement in F1 score over other methods was observed at lineage-level resolution for *C. acnes*. In contrast, PHLAME achieved relatively similar F1 scores to other methods when classifying *S. epidermidis* at the phylogroup level with a 25% held-out database. However, we note that PHLAME was able to estimate relative abundance profiles of *S. epidermidis* at the phylogroup level significantly more accurately than other methods at coverages below 5X (Fig. S7).

### PHLAME limits spurious detections of lineages

We additionally benchmarked PHLAME using a real microbiome sequencing dataset^19^ for which thousands of cultured isolate genomes and paired metagenomes were obtained from the same samples (Fig. 4A; Methods). This dataset was obtained from the facial skin of parents and children attending the same school and included isolates from two species: *C. acnes* and *S. epidermidis*. On average, each subject harbored seven lineages of each species; analysis of isolate data showed that subjects harbored both unique lineages and lineages shared with family members, but cross-family lineage sharing was very rare^19^.

**Figure 4:**
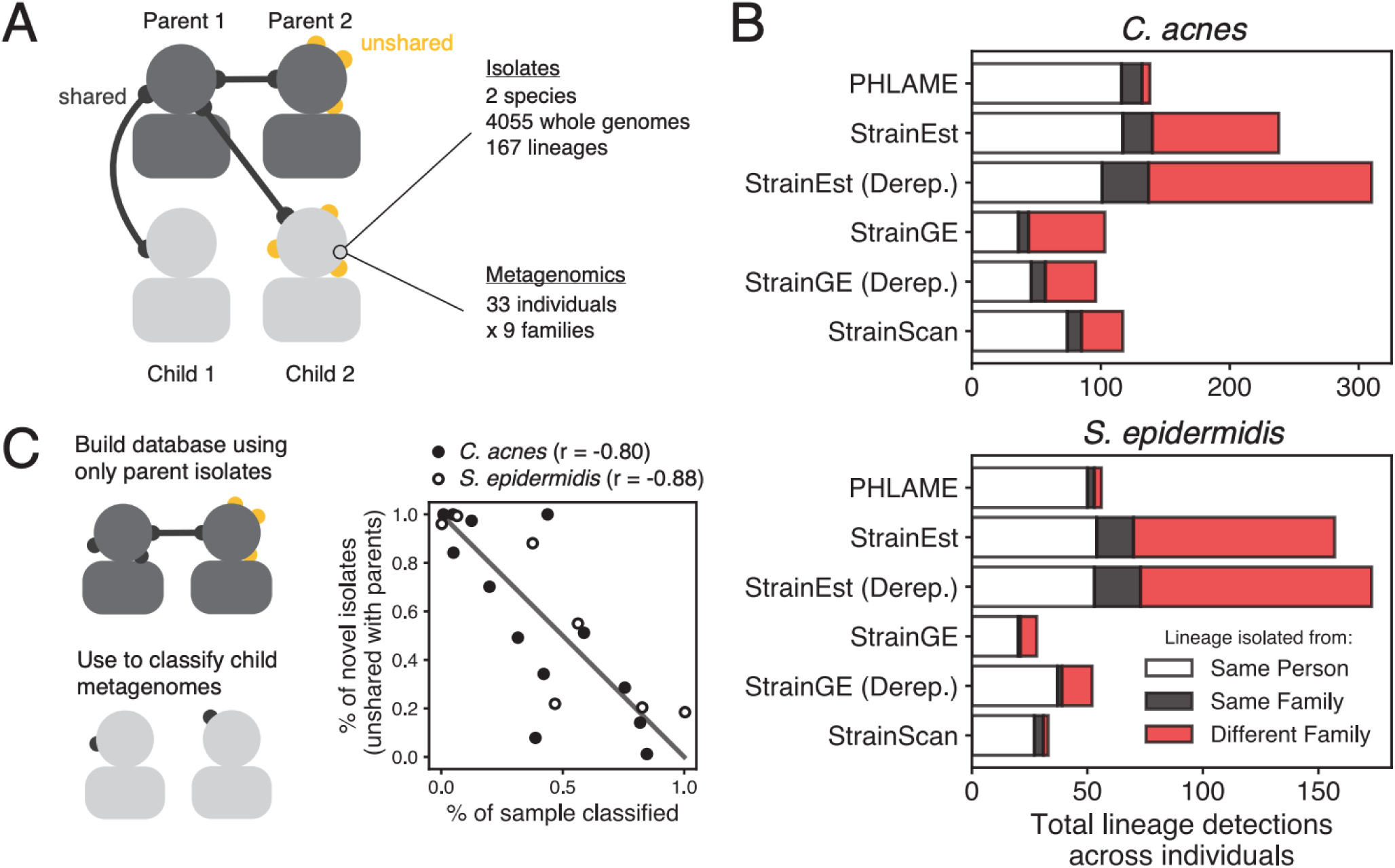
PHLAME limits spurious detections of lineages. **(A)** 4,055 isolate genomes and metagenomes obtained from the same skin swabs provide a unique opportunity to benchmark strain-resolved metagenomics methods using real microbiome sequencing data. **(B)** Total number of lineage detections from metagenomics across all individuals, colored by whether the lineage was originally isolated from the same person, the same family, or a different family. Lineages isolated from both the same person and same family are labeled as being from the same person. For methods that recommend dereplicating reference genomes before constructing a database, databases were constructed using either all the reference genomes or a single representative genome per lineage (Derep.) **(C)** PHLAME identifies samples that are abundant in novel lineage diversity. A database was constructed using only isolates originally cultured from parents and used to classify child metagenomes. For each child with metagenomic coverage > 1X for a given species, the percent of strain diversity that was classifiable via PHLAME is plotted against the percent of novel isolates (unshared with parents) that were obtained from t at child. Pearson’s correlation coefficients are shown next to each species.

We tested whether methods could identify lineages shared within families while not falsely detecting extensive cross-family lineage sharing. We did not expect the culturing data to represent the complete ground truth of individual strain diversity, nevertheless we counted metagenomic lineage detections on subjects as presumable false positives if the lineage was never isolated from that subject or any family members (Methods). PHLAME’s false positive rate was 4.4% *C. acnes* and 5.4% for *S. epidermidis*, while other methods reported a higher percentage of presumably false positive detections (41%, 55% for StrainEst, 40%, 25% for StrainGE, 27%, 6.1% for StrainScan; *C. acnes*, *S. epidermidis*) (Fig. 4B). Notably, PHLAME maintained high recall of true unique and within-family shared lineages, behind only StrainEst which also detected a very high percentage of false positives.

We next tested if PHLAME can identify samples that contain novel diversity at high abundances. We generated a reference database using only the genomes originally isolated from parent samples and used it to classify metagenomes obtained from child samples. If a lineage cultured from a child was not cultured from either parent, we considered it “novel” with regards to the parent-only database. We compared the proportion of “novel” isolates from each child to the proportion of the child’s metagenome that was classified at the lineage level using the parent-only database. (Fig. 4C). We recovered strong negative correlations between these two values for both species (r = −0.80 for *C. acnes*, r = −0.88 for *S. epidermidis*), indicating that PHLAME can identify samples that abundant in novel strain diversity for targeted culturing efforts.

### *Cutibacterium acnes* clades associate with human demographic features

We used PHLAME to search for intraspecies associations in *C. acnes* using healthy facial skin metagenomes obtained from 969 individuals and 3 geographic regions (USA, Europe, and China). Intraspecies diversity was classified using a reference database of 360 representative *C. acnes* isolates (Fig. 5A). We characterized diversity at a set of clades (phylogroups) that best matched an existing typing scheme for *C. acnes*^25^. Some phylogroups had their own distinct phylogenetic substructure, which we refer to as sub-phylogroups. We therefore profiled the presence and abundance of each phylogroup and sub-phylogroup in each sample. At the phylogroup level, novel *C. acnes* diversity was uncommon, and 92% of samples above 1X coverage were over 80% classified by PHLAME at phylogroup resolution (Fig. S13).

**Figure 5:**
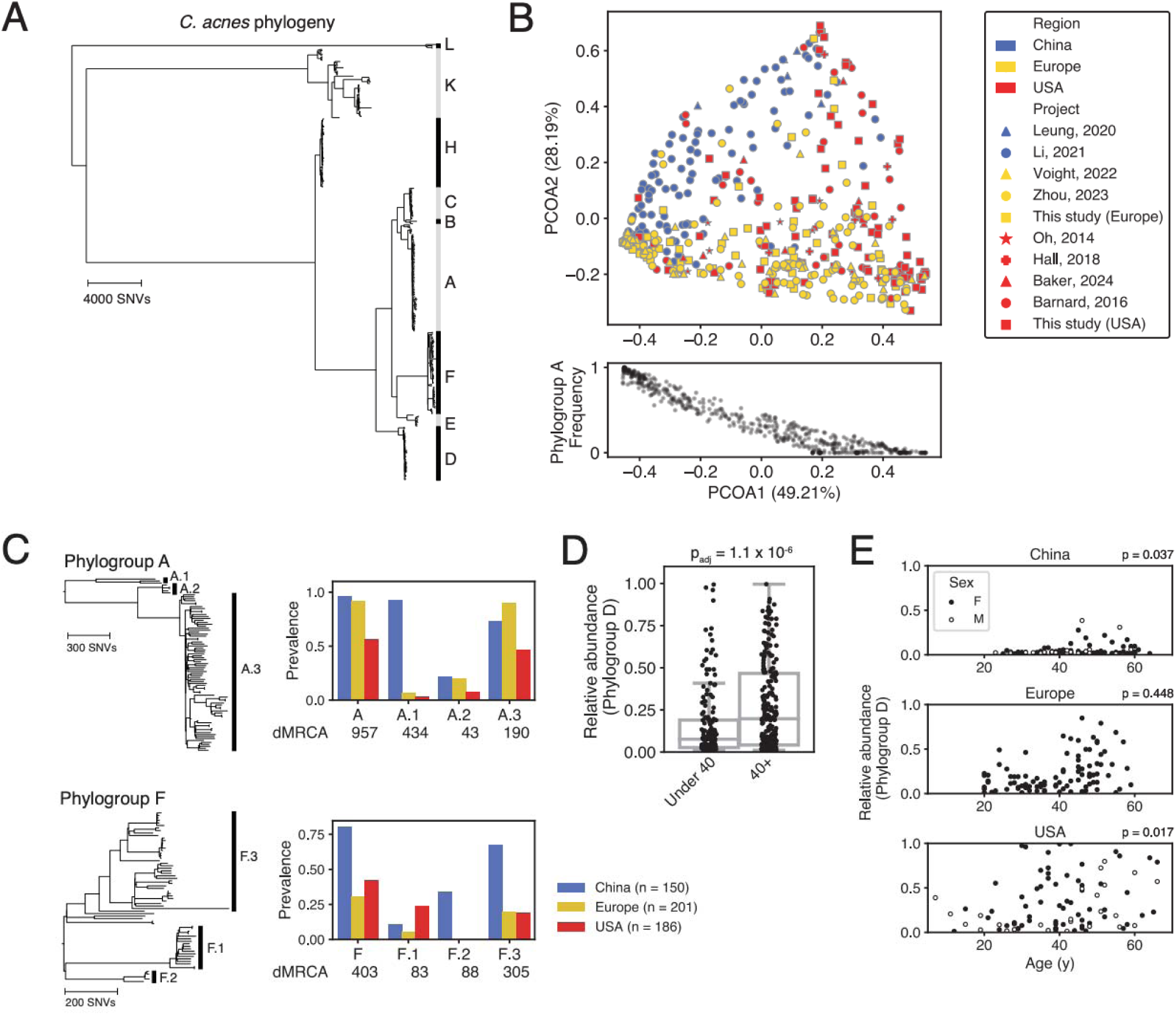
*C. acnes* clades associate with human demographic features. **(A)** Core-genome reference phylogeny of *C. acnes* constructed from 360 representative genomes. Major intraspecies clades (phylogroups) are indicated with letters. **(B)** On-person *C. acnes* diversity is regionally distinct. PCoA (Bray-Curtis dissimilarity) of *C. acnes* phylogroup abundances from 537 healthy subjects between the age of 18 and 40 that were over 80% classified at the phylogroup level by PHLAME. The major axis of variation between individuals (PCOA1, 49.21%) is correlated with the frequency of *C. acnes* phylogroup A (Spearman r = −0.95). **(C)** Some sub-phylogroups are distributed differently across regions from their parent phylogroup. Left: zoomed in view of the core-genome phylogeny for phylogroups A and F. Intra-phylogroup clades, called sub-phylogroups, are indicated with numbers. Right: Prevalence by geographic region for phylogroups A, F, and their corresponding sub-phylogroups. The dMRCA of each clade is shown below. **(D)** *C. acnes* phylogroup D achieves higher relative abundance on older individuals. Bar plot showing the difference in phylogroup D frequency on individuals under 40 compared to 40+. Adjusted p-value represents the result of a rank sum test after Bonferroni correction. **(E)** Full plot of the relationship between *C. acnes* phylogroup D abundance and age, broken down by geographic region and reported sex. Spearman rank correlation p-values for both sexes combined shown next to each geographic region.

We observed significant heterogeneity between on-person *C. acnes* populations from different regions. At the phylogroup level and among samples that were over 80% classified, *C. acnes* diversity clustered significantly by geographic region (PERMANOVA p = 0.001) (Fig. 5B, Fig. S11). The first major axis of variation between individuals (49.21% of variation) was strongly correlated with the on-person abundance of phylogroup A (Spearman r = −0.95). At the sub-phylogroup level, we observed two striking cases of near-complete geographic restriction to China (Fig. 5C). Sub-phylogroups A.1 and F.2 reached high prevalence across individuals within China (93% and 34% of individuals, respectively), and were exceedingly rare on individuals in our cohort sampled outside of China (5% and 0%, respectively, p<0.001 for both, Fisher’s exact test). Interestingly, these patterns appear incompatible with drift or co-migration with humans, as both sub-phylogroups possess extremely recent common ancestors (dMRCAs of 434 and 88 SNVs, respectively), and have sister clades with similar ages but cosmopolitan distributions (Fig. 5C). The high local fitness but limited spread of A.1 and F.2 may indicate the presence of selective pressures that restrict their success outside their region of origin.

We additionally tested whether certain clades of *C. acnes* were associated with age, as there is a well-characterized change in skin physiology on older individuals^27–29^. The median age for an adult in our cohort was 40; we therefore searched for differences in phylogroup and sub-phylogroup relative abundance on individuals under 40 compared to over 40 (Fig. S12A). We found that phylogroup D reached significantly higher relative abundance on individuals over 40 (p_adj_ = 1.1e-6). We were able to independently recover a significant association between phylogroup D abundance and age for both reported sexes and for two out of three regions (Fig. 5D, Fig. S12B-C). For the USA, this relationship was not significant between individuals under 40 compared to 40+, but there was significant rank correlation between age and abundance (p = 0.017, Fig. 5E). Together, these results suggest there is a largely generalizable association between host age and *C. acnes* phylogroup D abundance.

### *Gardnerella* diversity in the vaginal microbiome

To investigate intraspecies associations at a different body site, in a disease context, and in a taxon with a low overall number of reference genomes, we used PHLAME to profile the genus *Gardnerella*, a natural member of the human vaginal microbiome. Individuals with *Gardnerella*-dominant community types are at increased risk of adverse health outcomes, including bacterial vaginosis, preterm birth, and HIV^2,30^. Initially classified into a single species (*G. vaginalis*), it is now understood that the *Gardnerella* clade consists of many members distant enough to be separate species. Comprehensive analysis of the *Gardnerella* taxonomy is limited by poor sampling – currently, 7/13 proposed species have fewer than three representatives^31^. We reasoned that PHLAME’s novelty-aware approach would enable us to search for associations among the *Gardnerella* even with a limited reference database.

We built a reference collection of 87 publicly available *Gardnerella* genomes; because of the diversity of the clade, these were aligned to four separate reference genomes (Fig. 6A, Methods). We excluded four of the 13 proposed *Gardnerella* species from the database due to being either singletons or not aligning well to any of the four reference genomes. We used this reference database to classify 775 public vaginal metagenomes from four independent studies and 268 individuals. Samples with detectable *Gardnerella* usually contained many coexisting taxa (mean 3.9 ± 1.7; Fig. S14). Compared to *C. acnes*, less of the *Gardnerella* population in each sample could be confidently assigned to a known clade (Fig. S10). This lower assignment rate (two-sample K-S test p<0.001) mirrors the lack of available reference genomes and validates the novelty-aware behavior expected from PHLAME.

**Figure 6:**
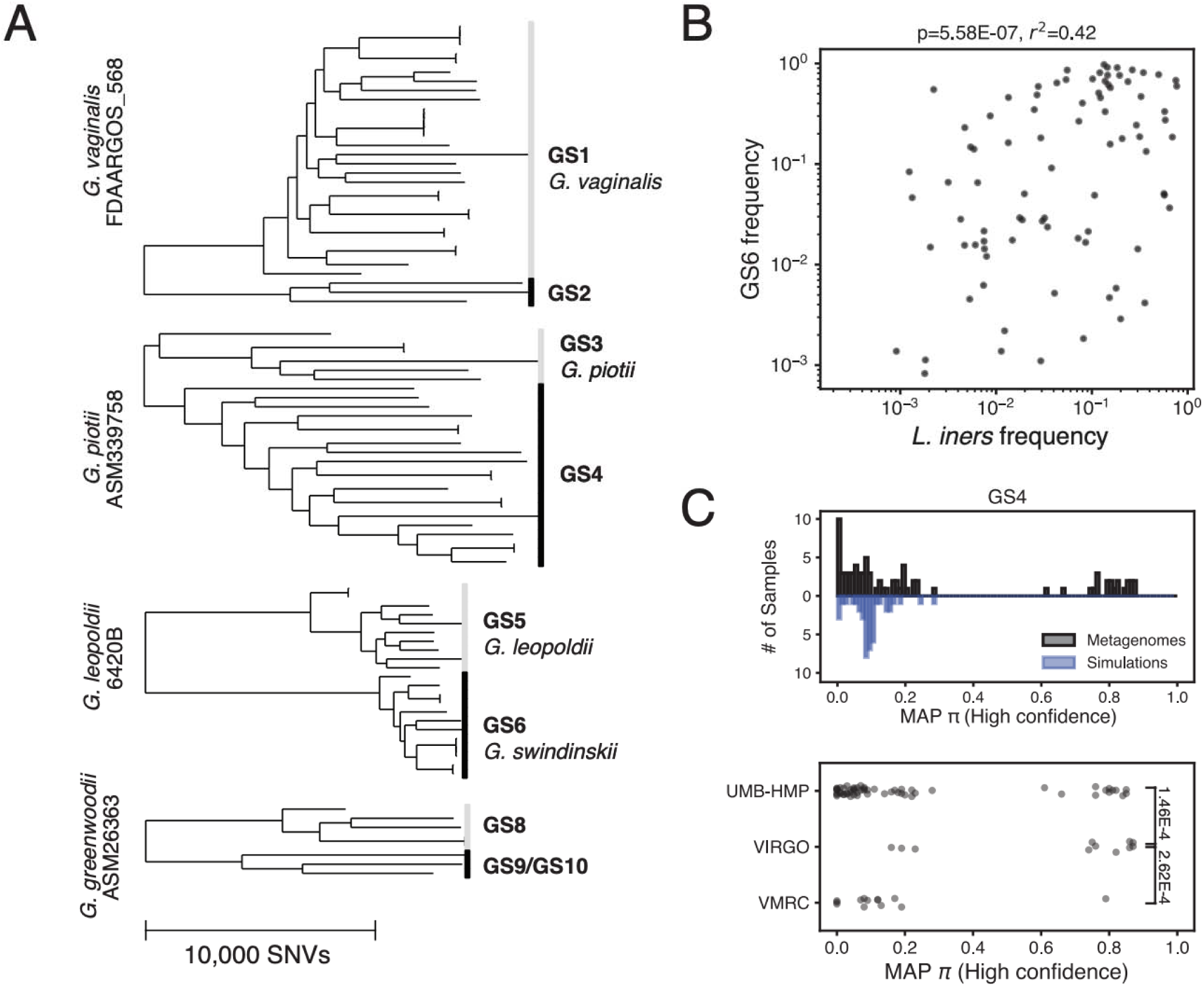
*Gardnerella* diversity in the vaginal microbiome. (A) Core-genome phylogenies of the *Gardnerella* taxon from 87 representative genomes, labeled with both proposed (GS prefix) and named species. Because of the diversity of the taxon, genomes were aligned to 4 reference genomes, which are listed before each tree. The scale bar shown is the same for all phylogenies. (B) Frequency of the *Gardnerella* clade GS6 within the overall *Gardnerella* population correlates with the frequency of *L. iners* in the sample. Spearman correlation coefficient and associated p-value are shown. (C) Some, but not all *Gardnerella* clades show evidence of related novel diversity. Top: High-confidence MAP estimates for π along clade GS4 from real metagenomes (black) and synthetic metagenomes comprised of random combinations of *Gardnerella* genomes (blue). π estimates from real metagenomic samples are differently distributed from synthetic metagenomes (p_adj_ = 0.01, two-sample KS test), with only the real metagenomic samples having a peak in π estimates near 0.8, suggesting the presence of a putative novel *Gardnerella* taxon. Bottom: MAP estimates for π along clade GS4, separated by study. Each dot represents one estimate from one sample. Samples from the VIRGO project are enriched in high π values compared to those form the UMB-HMP and VMRC project (Rank sum test, p-values shown in brackets).

We tested whether there was a significant difference in *Gardnerella* strain composition in communities with different species-level compositions. Previous studies have shown that *Gardnerella* are common in two vaginal community types: a *Gardnerella*-dominated community type and a *Lactobacillus iners*-dominated community type^2^. We chose 149 samples previously labeled as being either *L. iners*-dominated or *Gardnerella*-dominated from their species-level abundances (one sample per subject, Methods). We found that the percentage of *Gardnerella* assigned as clade GS6 was - positively associated with *L. iners* frequency in the community (Fig. 6B; Fig S15A). This association was not confounded by sequencing depth, as there was no relationship between *L. iners* frequency and the number of reads that mapped to the reference genome used for GS6 (Fig. S15B), and we were able to recover this relationship across multiple independent sequencing efforts (Fig. S15C). These results suggest a possible interaction between *L. iners* and *Gardnerella* clade GS6 in the vaginal microbiome, which may influence overall vaginal community composition and genital health.

Finally, we used PHLAME to infer the presence of novel *Gardnerella* diversity. We reasoned that high values of DV_b_ inferred across many samples may indicate the presence of putative novel clades. For each clade, we aggregated all high-confidence (highest posterior density interval < 0.2) estimates for π, regardless of whether the clade was counted as detected or not (Methods). While π estimates for these high-confidence calls were generally low, we observed that many clades had additional peaks at π values greater than 0 (Fig. 6C, Fig. S16). To ensure these peaks were not artefacts of known but rare *Gardnerella* species not included in our database, we created synthetic metagenomes comprised of random combinations of *Gardnerella* genomes. We then compared π estimates obtained from real samples to those obtained from synthetic metagenomes. Only 1% of π estimates from synthetic metagenomes were higher than our default detection threshold of 0.35, compared to 30% of estimates in real samples (Fig. S16, p_adj_<0.02 for all clades, K-S test). We highlight one example where a putative novel clade is inferred to share ancestry with clade GS4 and is relatively enriched in the VIRGO study (Fig. 6C; Fig. S17). Samples that were inferred to have substantial abundances (>20% of the overall *Gardnerella* population) of at least one putative novel clade are listed in Supplemental Table 8 and could make promising targets for future culturing efforts.

## Discussion

Taxonomic novelty is a challenge for all classification methods that use reference databases, particularly at the intraspecies level where nearly all strains are expected to be novel to some extent. We have presented PHLAME, a reference-based strain profiler that explicitly uses evolutionary history to quantify intraspecies novelty in metagenomic samples at multiple levels of resolution. We have shown how our strategy leads to improved performance and interpretability in the presence of novel strains (Fig. 3-4), and demonstrated its utility by identifying novel associations in public metagenomes (Fig. 5-6).

Although strain-resolved reference databases were overlooked in early microbiome research due to limited reference collections, they now show significant promise as genome collections continually expand. New techniques continue to expand the number of bacterial species that can be cultured^32^; even the notoriously difficult epibiont TM7 from the oral microbiome has been successfully cultivated by multiple groups^32,33^. Beyond providing robust performance across sample types and sequencing depths, reference-based approaches like PHLAME also naturally lend themselves to experimental follow-up, as cultured representatives of reference genomes are often readily available. Moreover, PHLAME’s ability to reveal samples with unexplored strain diversity (Fig. 4C, Fig. 6C) promises to establish a cyclic reference-based discovery pipeline, in which identification of unexplored microbial diversity guides targeted cultivation efforts that in turn improve intraspecies reference databases.

Using facial skin metagenomes from nearly a thousand people, we identified two recently differentiated clades of *C. acnes* that were highly enriched in China compared to Europe and the United States (A.1 and F.2; Fig. 5C). These observations are difficult to explain as co-diversification along with humans^34^, as both A.1 and F.2 possess such recent common ancestors that no plausible bacterial molecular clock rate^21,35^ could align their emergence with early human migration events (40,000 to 100,000 years ago^36^). The combination of high prevalence in China, near-complete geographic restriction to China, and the cosmopolitan distribution of closely-related sister clades of similar ages suggests that subphylogroups A.1 and F.2 have a recent origin either within or near China and are prevented from spreading globally by a selective barrier to migration. Further work characterizing genomic and phenotypic differences in these clades might reveal region-specific selective pressures in *C. acnes* and their corresponding adaptations.

We report a significant shift in *C. acnes* strain composition beginning in mid-life, driven by a robust increase in the relative abundance of *C. acnes* phylogroup D across reported sex and geographic region (Fig. 5D-E). While a previous study^37^ described a similar abundance shift between pre- and post-menopausal women, our results suggest this change is not specific to female menopause. Aging skin undergoes several sex-neutral changes in physiology, including epidermal thinning, lower cell turnover, altered immune activity^27^ and decreased surface water and lipid content^28,29^, all of which could support a selective change in strain composition. Importantly, future studies investigating the relationships of specific *C. acnes* strains with disease or other features should consider stratifying by age to account for this now-known variation in strain composition.

In the vaginal microbiome, we find a positive association between the proposed species *G. swidsinskii* (clade GS6) and coexisting *Lactobacillus iners* (Fig. 6B). Vaginal community composition has been extensively linked to female reproductive health, and studies have shown that *L. iners*- dominated vaginal microbiomes are more likely to transition into *Gardnerella*-dominated ones over time^38^ and raise the risk of adverse health outcomes^2,30^. Our results indicate that *L. iners*-dominated communities are also associated with a specific *Gardnerella* species, *G. swidsinskii,* but further investigation is needed to decipher the relevance of this taxon on community dynamics and human health. *G. swidsinskii* could be an important cross-feeder for *L. iners*, which is known to have key auxotrophies^38^ and a reduced genome compared to other *Lactobacilli*. Alternatively, it may act as an early invader of *L. iners*-dominant microbiomes, destabilizing the community for colonization by other *Gardnerella* taxa.

To guide appropriate interpretation of PHLAME results, we address a few important practical constraints. PHLAME currently makes a single prediction of DV_b_ per branch of the phylogeny, meaning that in scenarios where both a known and closely related novel clade are present in the same sample, the more abundant clade will dominate the output. In addition, DV_b_ may be difficult to interpret in terms of evolutionary time for bacteria with high rates of recombination, although we note that even highly recombinant species have significant population structure with clade-specific alleles^39,40^. Lastly, PHLAME currently only investigates intraspecies diversity on a single-species basis; further work will be required to make scale to whole microbiome analyses spanning many species.

PHLAME will help leverage accumulating metagenomic data to its fullest extent, especially in environments where high intraspecies diversity is the norm for most species, including human skin^19^ and the female genital tract^41^, aquatic ecosystems^16,42^, and soils^16^. Future studies using PHLAME will, classify intraspecies diversity at the level needed detect strain transmissions, guide targeted culturing efforts for novel intraspecies diversity, and reveal compositional strain-level associations with disease, behavior, and biogeography. Finally, PHLAME will inspire other approaches for novelty quantification in reference-based metagenomic interpretation.

## Methods

PHLAME is a complete pipeline for the creation of intraspecies reference databases and the metagenomic detection of intraspecies clades, their relative frequency, and their estimated divergence from the reference phylogeny. The accepted raw input(s) to PHLAME are: [1] a species-specific assembled reference genome in fasta format [2]. A collection of whole genome sequences of the same species in fastq or aligned bam format, and [3] metagenomic sequencing data in either fastq or aligned bam format.

### SNV calling

In the first step, whole genome sequences (reference sequences) are aligned to a species-specific reference genome and variants in each genome are identified. Raw fastq files are aligned using bowtie2 (v.2.2.6 -X 2000 --no-mixed --dovetail) and nucleotide counts across each position are collated using SAMtools mpileup (v.1.5 -q30 -x -s -O -d3000). Nucleotide counts are compiled into a data structure and base calls are determined by taking the major allele nucleotide at each position, for each sample. In order to distinguish high-confidence base calls from low-confidence ones introduced by low coverage, sequencing errors, alignment errors, or non-pure samples, a set of filters are applied to genomes, positions, and base calls within a genome. First, genomes are excluded if the median coverage across the sample is below 8X. Base calls are marked as ambiguous (“N”) if the FQ score produced by SAMtools is above −30, the coverage per strand is below 3X, the major allele frequency is below 0.85, or more than 33% of reads supported an indel within 3 base pairs up or downstream of the position. Positions are then discarded from the alignment if greater than 10% of the samples support an N at that position, if the median coverage across samples for that position is below 5X, or if the median coverage across samples for that position is more than 2 times greater than the median coverage across samples for all core positions (a proxy for copy number). Finally, genomes are additionally excluded if greater than 10% percent of post-filtering positions in that sample are considered ambiguous.

### Phylogenetic tree construction and clade definitions

Positions containing at least one polymorphism after filtering are used to generate a reference phylogeny. PHLAME packages RAxML (v.8.2.12 -m GTRCAT) for convenience but will accept common tree file formats (Newick, NEXUS) produced by any available phylogenetic reconstruction method. Where possible, PHLAME will scale tree branch lengths into core-genome SNV distances when the Pearson correlation coefficient between pairwise branch lengths and core-genome SNV distances (calculated as the Hamming distance between base calls) is > 0.75.

PHLAME generates hierarchical intraspecies typing schemes from phylogenies by first defining candidate clades, then searching for identifiable SNV barcodes for each candidate clade. PHLAME will consider any clade that meets a set of ‘minimum identifiable’ criteria. This allows PHLAME to search for phylogenetic associations across any subset of intraspecies diversity. Default criteria used to include a candidate clade are a minimum of 4 genomes in the clade, a branch length of at least 1000 SNVs, and a bootstrap value > 0.75.

### Clade-specific SNVs

PHLAME searches for a set of clade-specific SNVs for each candidate clade by looking for the set of alleles that [1] are common to all members of a clade that have a non-ambiguous base call at that position, [2] are ambiguous in less than 10% of in-clade members, and [3] are not found in any other clades. To account for scenarios where reads may map to several closely-related taxa, users can optionally include a collection of outgroup genomes assumed to competitively recruit reads from the species of interest. Positions that have a non-ambiguous base call in greater than 10% of a set of user-defined outgroup genomes are additionally discarded. Candidate clades with fewer than a minimum number of clade-specific SNVs (default 10) are removed from consideration.

### Metagenome classification

Metagenomic sequences are first aligned to the same species-specific reference genome used to build the database (bowtie2 v.2.2.6 -X 2000 --no-mixed --dovetail) and nucleotide counts across each position are collated using SAMtools mpileup (v.1.5 -q30 -x -s -O -d3000). Nucleotide counts across the informative positions containing a clade-specific SNV are used to infer the clade composition in a sample. By default, PHLAME will only proceed with DV_b_ inference and abundance estimation if more than 10 positions have at least one read supporting the clade-specific allele

The number of reads containing a clade-specific SNV (x_i_) at position *i* is assumed to be generated from the following process:

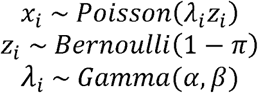

Parameters for the prior distribution over α (a and b) are determined from the total number of reads spanning all alleles at the same positions *y_i_*, where 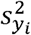 is the sample variance of *y_i_*.

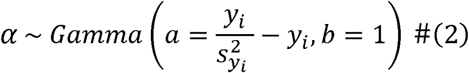

We use a Slice-within-Gibbs sampling scheme to estimate the posterior distribution for the parameters α, λ, and π (see Supplemental Methods). Chains by default are run for 10,000 steps with the first 10% discarded as burn-in. In order for a clade to count as present, 50% of the posterior density over π must be below a defined threshold (default 0.35) and the lower bound of the 95% highest posterior density interval over π must be less than 0.1. These default thresholds were chosen based on performance in real-data benchmarking (Fig. S9).

The relative abundance of each detected clade is calculated as the ratio between the maximum a posteriori (MAP) estimate for the rate parameter across clade-specific alleles (λ_clade-specific_) and the MAP estimate for the rate parameter across all alleles at the same positions (λ_all_), which is assumed to originate from an ordinary negative binomial distribution.

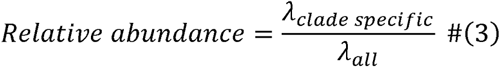

Because the relative abundance of each clade is determined independently, the total frequency of a set of non-overlapping clades may sum to below 1, or rarely above 1. Total frequencies that sum to below 1 inform what proportion of the sample is composed of novel clades, but total frequencies that sum above 1 are obviously not possible. We recommend that samples with total frequencies that sum above 1 are normalized down to 1, as we do in all analyses described.

### Novel diversity benchmarking

We characterized the behavior of three reference-based strain classification methods: StrainEst, StrainGE, and StrainScan in the presence of novel diversity by masking individual branches from the reference set and evaluating performance when faced with a genome from the held-out clade.

Following the procedure laid out in <u>Phylogenetic tree construction</u>, core-genome phylogenies were generated for 3 species: *C. acnes, S. epidermidis*, and *E. coli*, and clades were defined for each phylogeny (Fig S2, Supplemental Tables 4-6). Whole genome sequences for *C. acnes* and *S. epidermidis* were obtained from two studies^18,19^. For *C. acnes*, we also included genomes labeled as ‘Cutibacterium acnes’ in the 661K bacterial genome database^43^, as well as four *C. acnes* isolates (Cacnes_PMH5, Cacnes_JCM_18919, Cacnes_JCM_18909, Cacnes_PMH7) with whole-genome assemblies but no raw sequencing data representing phylogroup L. We used wgsim (v.0.3.1; -e 0.0 -d 500 -N 500000 -1 150 -2 150 -r 0.0 -R 0.0 -X 0.0) to generate synthetic raw sequencing data from genomes where only the assembly was available. For *E. coli*, whole genome sequences were obtained from the ECOR collection^44^ and an existing study containing 976 sequenced *E. coli* isolates^45^. Genomes were dereplicated as follows: for the Conwill et al.^18^, Baker et al.^19^, and Thänert et al.^45^ datasets, a single representative isolate was chosen per lineage (as defined in each study) based on highest median coverage. All other genomes were dereplicated by first clustering based on core genome SNV distances using dbscan (epsilon = 500 SNVs, minimum number of genomes in a cluster = 2), then choosing a representative isolate from each dbscan cluster based on highest median coverage, as well as all singletons.

Genomes were held out from databases as follows: for each node with a bootstrap value > 0.75, all genomes descended from one of the daughter branches were held out from the reference set, and a new reference database was built for each method. This was repeated for every possible combination of well-supported nodes and branches. We did not include scenarios where one of the two branches originating from the root was held out to avoid artefacts driven by midpoint rooting of the phylogeny. Sequencing data from one held-out genome was subsampled to 10X and classified against each method’s reference database. We used the Hamming distance between core-genome base calls to measure genetic distance between pairs of genomes. We evaluated each method by looking at the difference in distance between the output genome(s) and the held-out genome (Δ_i_) compared to the distance between the true closest genome in the database and the held-out genome (Δ_t_). Because methods often output multiple genomes, we normalized each distance by the vector of relative abundances reported in that sample (R_i_ = R_1_ … R_Ngenomes_).

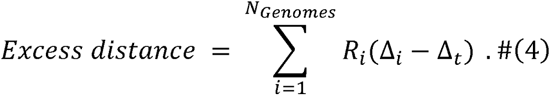

The same hold-out simulations were used to check the accuracy of divergence estimates given by PHLAME. We iteratively held out all descendants of a single branch from the PHLAME reference database, once again excluding instances where one of the two branches originating from the root was held out. We evaluated PHLAME’s ability to accurately measure the divergence of a genome belonging to the held-out clade at coverages ranging from 10X to 0.1X. The maximum a posteriori estimate of π was compared against the true divergence of the novel strain, defined as in (1) (Fig. 2E, Fig. S4).

### Synthetic metagenome benchmarks

Synthetic metagenomes were generated by combining raw reads from real whole genome sequencing data. In each metagenome, reads from five random sequenced genomes of either *C. acnes* or *S. epidermidis* were combined at random lognormal abundances (µ = 1, σ = 1), along with nine genomes from other species at constant abundances (Supplemental Table 1). For *C. acnes* and *S. epidermidis*, additional mock community members consisted of genomes from known human commensal skin bacteria (Supplemental Table 1).

We compared performance at a coarse-grained phylogenetic resolution (phylogroup level) and a fine-grained phylogenetic resolution (lineage level). Phylogroups were determined using existing typing schemes created for these species^26,46^; lineages are highly related clades that are separated by fewer than 100 mutations across the core genome^19^. We compared performance using both a perfect database, where all genomes in a sample are additionally found in the reference database as well as a database where 25% of the lineages or phylogroups were randomly held out (Fig. 3).

For StrainEst, StrainGE, and StrainScan, we compared performance using both a complete reference database and a database containing a single representative genome per lineage and chose the database with the best performance for each simulation in Fig. 3. To convert reported frequencies of individual reference genomes into clade frequencies, we summed the reported frequencies of every descendant of a given clade to obtain the total clade frequency. For PHLAME we chose the default parameters recommended; we show that results are consistent across reasonable classification parameters (Fig. S8). For all methods, we required the total clade frequency to be greater than 1% to be counted as detected.

### Real-data benchmarking

We additionally benchmarked the performance of each method (PHLAME, StrainEst, StrainGE, StrainScan) using a unique ground truth dataset consisting of thousands of isolates paired with metagenomes sequenced from the same samples. The Baker et al. dataset^19^ consists of 2,030 *C. acnes* and 2,025 *S. epidermidis* isolates along with paired metagenomes obtained from swabbing 33 subjects from 8 families. Metagenomic coverage for each species ranged from 1.1X-625X for *C. acnes* (median 51X) and 0.1X-76X for *S. epidermidis* (median 2.6X). For each species, a reference database was constructed using either the full isolate collection or a single representative genome from each lineage for methods that recommend dereplicating closely related isolates (StrainGE, StrainEst). We profiled metagenomes with each method under comparison, using the same classification parameters as in (Synthetic metagenome benchmarks). Reference genome frequencies were summed into clade frequencies as in (Synthetic metagenome benchmarks) and lineages were counted as detected if they reached a summed abundance above 1%. Metagenomic lineage detections were counted as presumable false positives if the family did not have isolated representatives of that lineage (rare instances of cross-family lineage sharing as determined from isolate data were counted as true positives). For both species, we only compared metagenomic data from families with at least ten cultured isolates of that species. We masked all detections of two *C. acnes* and two *S. epidermidis* lineages suspected of extensive cross-sample contamination (defined in the original publication^19^) from all methods.

To evaluate PHLAME’s ability to target samples with abundant novel diversity, we built a reference database using only genomes originally isolated from parents and used this database to classify metagenomic samples from children. We measured the total percent of each sample that PHLAME was able to classify at the lineage level. The percent of “novel” isolates on each child was measured as the number of isolates belonging to a “novel” lineage (i.e., unshared with parents) over the total number of isolates obtained from that child.

### Skin metagenome sample collection and sequencing

We collected and sequenced facial skin metagenomes from 443 individuals from Europe and the US in two independent efforts. Samples from the US were collected from healthy volunteers via self-swab with ESwabs (481C, Copan), stored in Amies buffer, and mailed overnight to Parallel Health Labs in San Francisco before processing immediately upon receipt. For DNA extraction, samples were subjected to selective lysis to remove human DNA by incubating with 0.04% saponin and 4 units of Turbo DNAse (AM2238, ThermoFisher) at 37°C with gentle shaking. Turbo DNAse was deactivated with 40 mM NaOH and 15 mM EDTA at 75°C for 15 min. Solution was neutralized with 80 mM Tris-HCl, pH 7 and 20 mM MgCl2. Microbial cells were spheroplasted with Metapolyzyme (MAC4L, Sigma Aldrich) at 35°C for one hour. Cells were further lysed with 1mg/mL Proteinase K and 0.5% SDS at 55°C for 30 min and finally mechanically lysed by beating with a 1:1 ratio of 0.1 and 0.5 mm glass beads (Biospec) in a Tissuelyser II for three minutes at 30 Hz. DNA extracts were purified with 1.2x CleanNGS SPRI beads (CNGS500, Bulldog Bio) and ethanol washes before resuspension in 10 mM Tris-HCl, pH 8. DNA libraries were generated with Illumina NexteraXT kits according to manufacturer instructions and sequenced on an Illumina NovaSeq 6000. Adapters were removed from metagenomic sequencing reads using cutadapt (v.1.18) and reads were filtered using sickle (v.1.33; -g -q 15 -l 50 -x -n) before aligning to the Cutibacterium acnes C1 reference genome. SAMtools markdup (v.1.15.1; -r -s -d 100 -m s) was used to remove duplicate reads from each sample. We report only the aligned .bam files as publicly available data.

Facial skin samples from Europe were collected from healthy volunteers via dry swab by the Skin Research Institute under an IRB approved protocol, placed in Amies buffer and immediately frozen. For DNA extraction, samples were thawed, then 250ul of buffer and the swab were transferred to the DNeasy PowerSoil HTP 96 kit. Sample DNA concentration was quantified using a SYBR Green fluorescence assay prior to library prep. After quantification, DNA isolated from swabs was normalized and used to generate libraries for Illumina sequencing using the published Hackflex protocol^47^. Briefly, DNA was fragmented using bead-linked tagmentation enzymes; these DNA fragments were then amplified and Nextera standard sequencing barcodes added using KAPA Mastermix PCR. SPRI beads were used for DNA purification and size selection. Samples were sequenced on an Illumina NovaSeq 6000, resulting in 150bp paired-end reads and approximately 100 million reads per sample. Reads were aligned to the human genome to discard host DNA sequences; only unaligned reads were used for further analysis.

### Metagenomic analysis of intraspecies variation in healthy facial skin

We searched the NCBI SRA database for publicly available human facial skin metagenomes using the keywords ‘skin metagenome’ and ‘human skin metagenome’. Studies were excluded if they had fewer than 10 subjects, did not include subject identifiers (to avoid duplication), or sampled subjects that had recently taken antibiotics or other skin therapeutics. We supplemented this public data with two newly-sequenced datasets collected from individuals from the USA and Europe. Subjects were removed from the dataset if they were diagnosed with acne or other skin diseases according to the metadata provided by each study. Samples were removed if they did not sample facial skin. Metagenomic sequencing data was dereplicated by subject as follows: for studies that collected samples at multiple timepoints per subject, the time point with the highest sequencing depth was chosen. For studies that collected samples at multiple facial sites per subject, the facial site with the highest average sequencing depth for each study was chosen, followed by the time point with the highest sequencing depth if applicable. For studies that collected skin microbiome samples using multiple sampling methods, we chose samples that were collected via facial swab to maximize method consistency with the rest of the dataset. In total, our combined dataset of facial skin metagenomes includes 969 individuals from 3 continents and 10 independent studies (Supplemental Table 2). Metagenomic sequencing reads were quality filtered using sickle (v.1.33; -g -q 15 -l 50 -x -n) and aligned to the Cutibacterium acnes C1 reference genome. SAMtools markdup (v.1.15.1; -r -s -d 100 -m s) was used to remove duplicate reads from each sample. Aligned sequencing reads were run through the PHLAME classifier using a custom-made *C. acnes* reference database with default classification parameters (-d 0.35 -p 0.5 -h 0.1). We used the *C. acnes* reference database built as described in Novel diversity benchmarking. In total, 360 representative isolates were included in the reference database. Whole genomes were taken through the PHLAME reference database creation pipeline with default parameters (-n 0.1 -p 0.1).

Clades were counted as detected if the estimated relative abundance was less than 1%. To generate a PCoA of variation in individual strain composition, we chose the non-overlapping set of clades (phylogroups) that corresponded to an established SLST scheme for *C. acnes*^46^. To control for potential confounders related to differences in strain composition with age, PCoAs and comparisons between regions were only done on the subjects in our dataset between the age of 18 and 40. Many subjects only had their approximate age reported (for example, “30s”); these subjects were included in analyses involving binary partitioning of ages (Fig. 5D, Fig. S12A-C) but not in continuous rank correlations (Fig. 5E).

### Metagenomic analysis of *Gardnerella* variation in the vaginal microbiome

We analyzed *Gardnerella* variation in 775 publicly available vaginal metagenomes obtained from four publicly available sequencing efforts^48,49^ (Supplemental Table 3). Metadata and vaginal community types were readily available for these studies in associated publications^48,49^. To generate a reference database for the *Gardnerella* taxon, we retrieved *Gardnerella* whole genome sequences from one existing study^41^ and downloaded additional genomes labeled as ‘Gardnerella vaginalis’, ‘Gardnerella piotii’, ‘Gardnerella leopoldii’ or ‘Gardnerella swidsinskii’ from the NCBI SRA and Assembly databases. We used wgsim (v.0.3.1; -e 0.0 -d 500 -N 500000 -1 150 -2 150 -r 0.0 -R 0.0 -X 0.0) to generate synthetic raw sequencing data when only the assembly was available. Because of the diversity of the clade, we made four reference databases each using a different reference genome, corresponding to the known cpn60 types of *Gardnerella*^50^. First, all Gardnerella genomes were aligned to each of the four reference genomes. Genomes were assigned to a given reference genome if less than 40% of positions in the alignment were marked as ambiguous (as defined in SNV calling). Genomes that did not fulfill this criterion for any reference genome or passed this criterion for multiple reference genomes were discarded. After filtering, a total of 87 *Gardnerella* genomes were included across our four databases. (Supplemental Table 7) Finally, for each of the four references, post-filtered genomes that were not assigned to that reference were labelled as outgroup genomes. Positions were excluded from a database if >10% of outgroup genomes had a non-ambiguous call at that position.

In order to search for associations between *Gardnerella* diversity and overall species abundances, we first identified metagenomic samples that were labeled as either community type III (*L. iners* dominated) or type IV (*Gardnerella* dominated) from existing metadata^49^. Filtered samples were dereplicated into 149 subjects by picking the sample with the highest sequencing depth per subject. Subjects that had less than 3X coverage across all reference genomes were discarded. We measured the species-level profile of each sample using kraken (v.2.0.9; default parameters) and bracken (v.2.6.0; default parameters). We measured the *Gardnerella* diversity in each sample using PHLAME with default classification parameters (-d 0.35 -p 0.5 -h 0.1)

To characterize novel diversity in the Gardnerella taxon, we collated MAP π estimates for each sample and each clade, regardless of whether the clade was reported as detected in the sample. We filtered for only high-confidence π estimates by filtering for only posterior distributions that had a highest posterior density interval less than 0.2. To check that peaks observed at high values of π were not artefactual, we generated 100 random synthetic metagenomes by combining reads from one genome for each defined *Gardnerella* clade, as well as one random genome from one of the four proposed *Gardnerella* species^31,41^ that were not included in our reference database (GS7, GS11, GS12, GS13). Genomes were subsampled at random lognormal abundances (µ = 1, σ = 1) to a total of 40X coverage across all 4 reference genomes.

## Author contributions

Conceptualization: EQ, TDL; Methodology: EQ, JSB, LM, VK, TDL; Investigation: EQ, JSB, ADT, CM, TDL; Funding acquisition: TDL; Writing: EQ, JSB, LM, VK, ADT, CM, TDL.

## Acknowledgements

We thank Nathan Brown, Ghislain Schyns, and Mathias Gempeler for subject recruitment and sampling of skin microbiomes. We thank Travis Gibson and Younhun Kim for assistance with the modeling and feedback on the manuscript. We thank Greg Fournier, Eric Alm, and members of the Lieberman Lab for helpful comments. This work was funded by a NSF GRFP fellowship to EQ, NIH grant 1DP2GM140922 to TDL, and a grant from DSM-Firmenich to TDL.

## Data availability statement

Sequence data from newly sequenced metagenomes from this study are available under BioProject PRJNA1211286. All publicly available genomes and metagenomes used in this study are listed in Supplementary Tables 2-7. PHLAME is an open-source Python package on GitHub (https://github.com/quevan/phlame). Data and code to recreate the findings of this paper are in the same GitHub repository.

## Supplemental Figures

**Figure S1:**
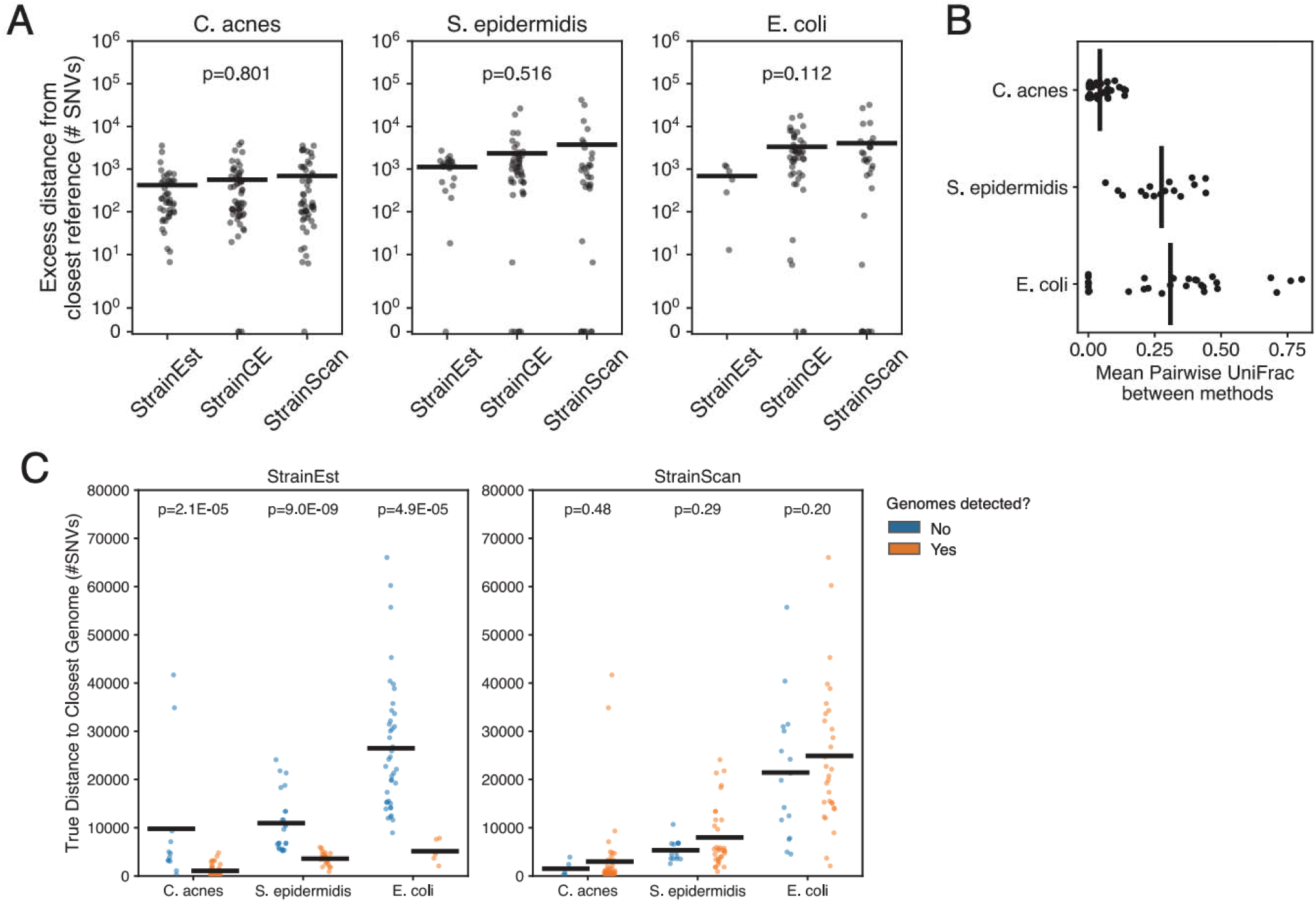
Behavior of strain-level classification algorithms when reference databases are systematically missing diversity. Results of three existing strain-level profiling methods when a reference database missing a section of phylogenetic diversity is used to classify a random held-out genome subsampled to 10X coverage across the reference genome. In the majority of simulations, the held-out genome is detected as one or more known genomes. (A) Excess Distance (Methods) between the true held-out genome and the detected genome(s), compared to the distance between the held-out genome and its true closest reference in the database. P-value shown is the result of a Kruskal-Wallis test for difference between methods. (B) Consistency in method outputs, as measured by the mean pairwise UniFrac distance between outputs of each method. For *E. coli*, this metric was only calculated between StrainGE and StrainScan, as StrainEst did not return an output for a majority of simulations. (C) More novel strains, as measured by the distance (# SNVs) to the closest reference genome, are less likely to be detected as a known genome by StrainEst, but not StrainScan or StrainGE (no graph shown for StrainGE as genomes were detected in all simulations).

**Figure S2:**
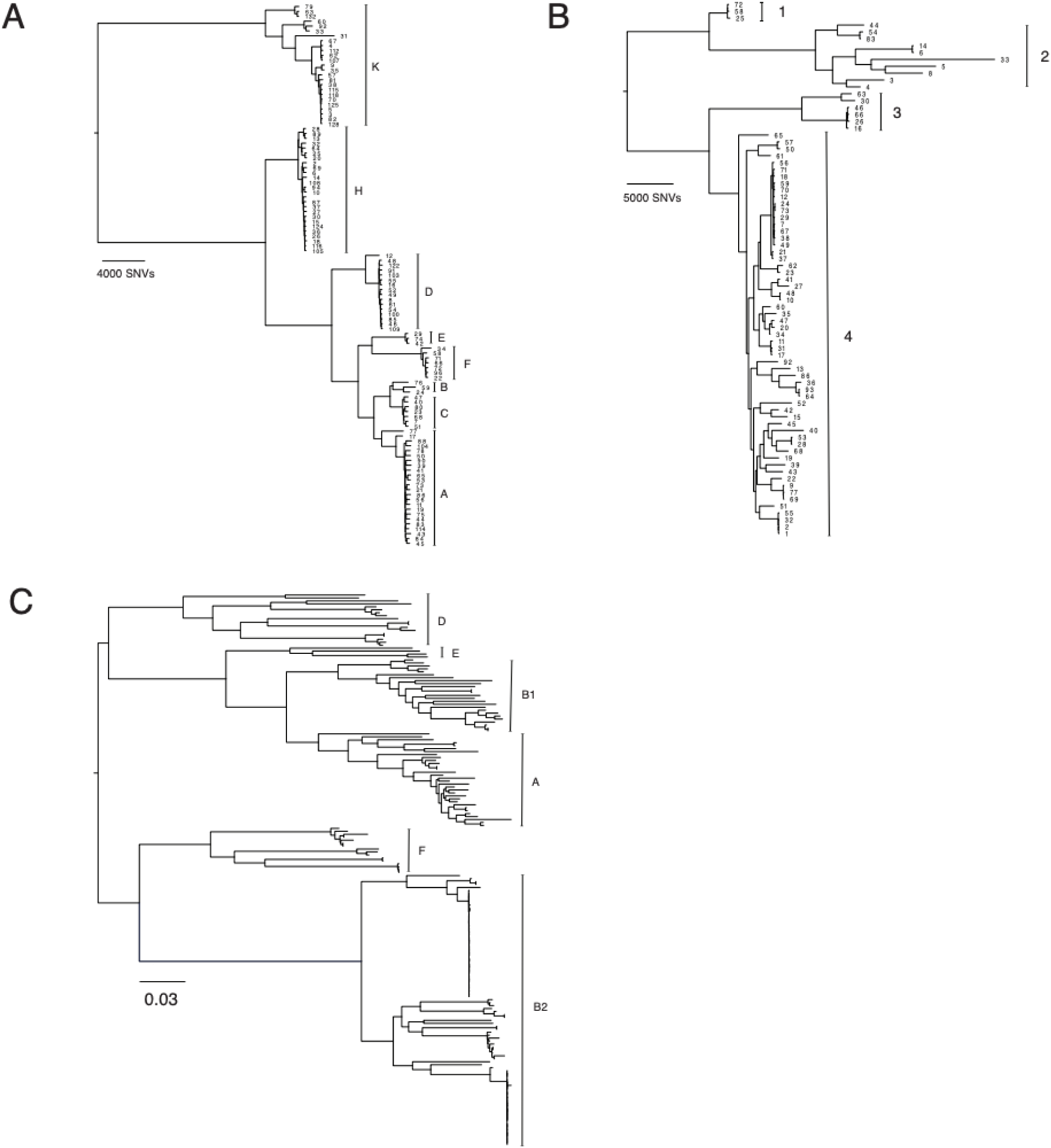
Phylogenies for *C. acnes*, *S. epidermidis, E. coli* used in held-out clade simulations. Core-genome maximum likelihood phylogenies (constructed using RaXML v.8.2.12) for (A) *C. acnes*, (B) *S. epidermidis,* and (C) *E. coli*. Lineages are labelled with a number, while phylogroups represent major intraspecies clades and are labelled with bars. Representative genomes were chosen for each lineage based on highest sequencing depth.

**Figure S3:**
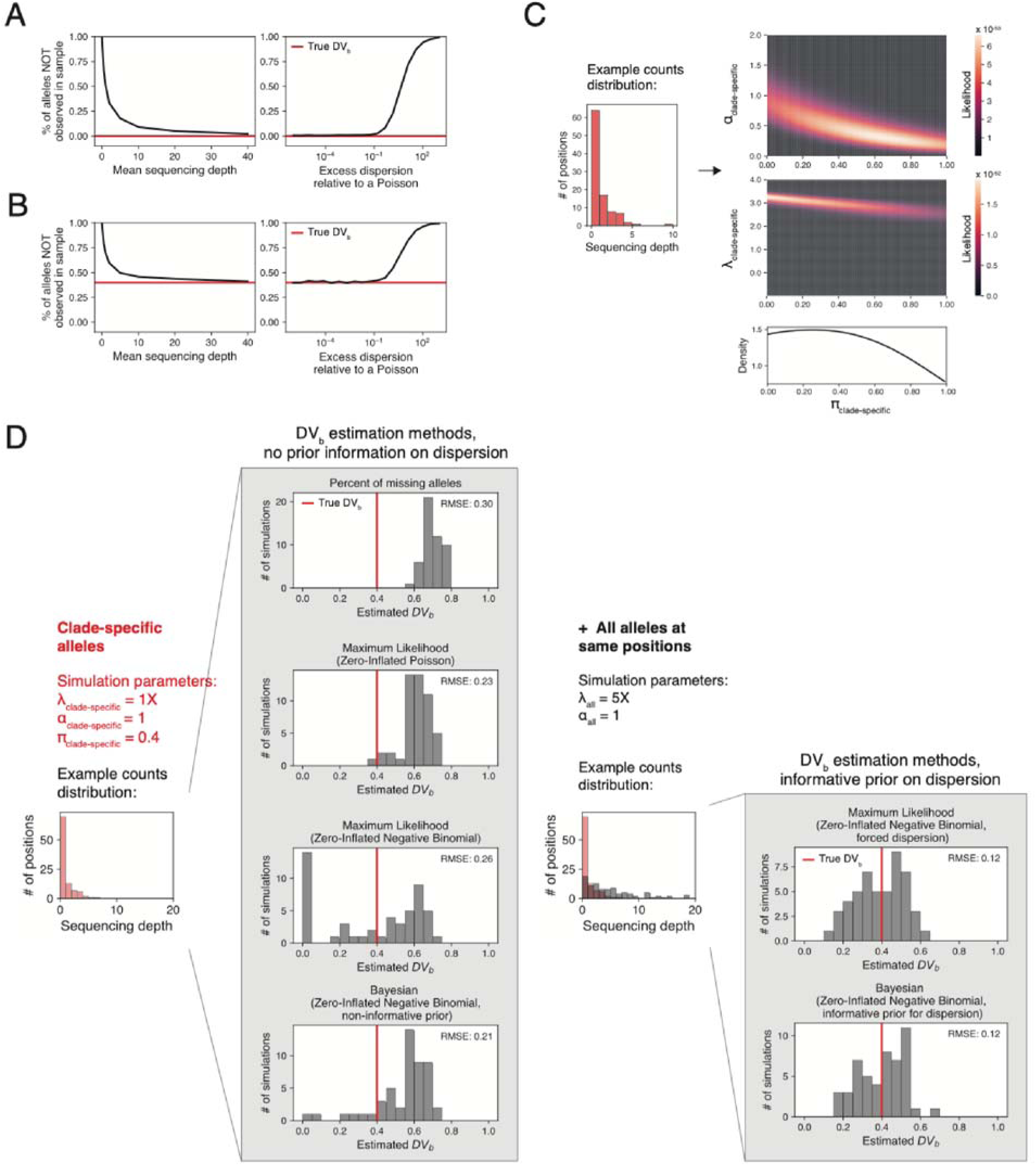
Prior information on read dispersion helps to constrain DV_b_ estimates. (A-B) The portion of missing clade-specific alleles does not necessarily reflect true values of DV_b_. We generated hypothetical read counts across a set of clade-specific alleles by simulating from a zero-inflated negative binomial (ZINB) distribution, varying either sequencing depth (left; from 0.1X to 40X) or overdispersion relative to a Poisson (right; from 10^-6^ to 10^3^). Values shown are the mean of 50 simulations per parameter set. The observed portion of missing clade-specific alleles begins to diverge from the true DV_b_ at sequencing depth <10X and overdispersion relative to a Poisson > 0.5. (C) When just using information on the number of clade-specific markers, a wide range of possible parameter combinations can reasonably explain any given counts distribution. Left: A random distribution of read counts supporting a set of clade-specific alleles. Right: Heatmap of the ZINB likelihood given the left distribution along parameters α_clade-specific_, π_clade-specific_, and λ_clade-specific_(see Fig. 2B). The marginal density of π_clade-specific_ (bottom right) supports a wide range of plausible values. (D) PHLAME overcomes this uncertainty by setting a prior on dispersion (represented by the parameter α_clade-specific_) using the coverage of all alleles at the same positions. Left: Attempts to measure the true DV_b_ value using without prior knowledge of the dispersion returns inaccurate results at low depth and realistic overdispersion. We simulated 50 hypothetical read count distributions from a ZINB distribution, then measured DV_b_ using the various methods, including the proportion of missing alleles, a maximum-likelihood zero-inflated Poisson and ZINB model, as well as a Bayesian ZINB model without an informative prior. The root mean squared error of each method is shown next to each plot. Right: Including prior information on the dispersion improves accuracy in the same simulation set. Here, we additionally simulated a read count distribution to represent the coverage of all alleles at the same set of positions as the clade-specific alleles. Including this information in either a maximum-likelihood or Bayesian inference algorithm improves RMSE compared to estimating DV_b_ from only the counts distribution across clade-specific alleles.

**Figure S4:**
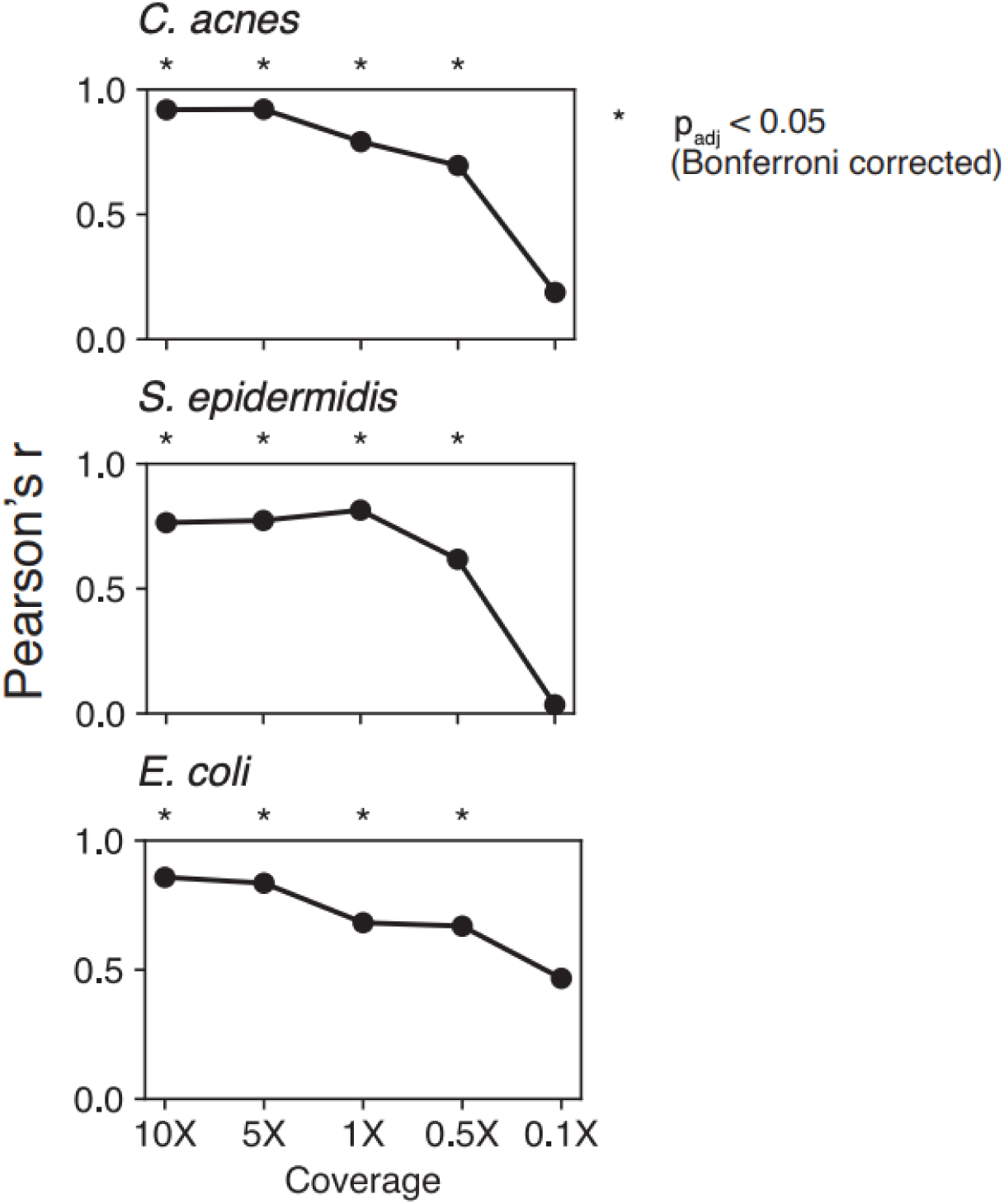
Accuracy of metagenomic estimates of DV_b_ with varying per-clade coverage. Pearson’s correlation coefficient between metagenomic point estimates of DV_b_ (represented by the parameter π) and ground truth DV_b_ values determined from the species phylogeny. Metagenomic inferences were obtained by constructing a reference database in which a single clade and all its descendants were held out, then using that database to classify a simulated metagenome containing a single genome form the held-out clade (See Fig. 2E). Simulated metagenomes were run through default PHLAME classification parameters, which requires minimum of 10 positions to have at least one read supporting the clade-specific allele. Low-coverage simulations that did not fulfill this criterion were not included in correlation calculations. While inference accuracy decreased at lower coverage, significant correlations (shown with stars) were recovered at per-clade coverages as low as 0.5X.

**Figure S5:**
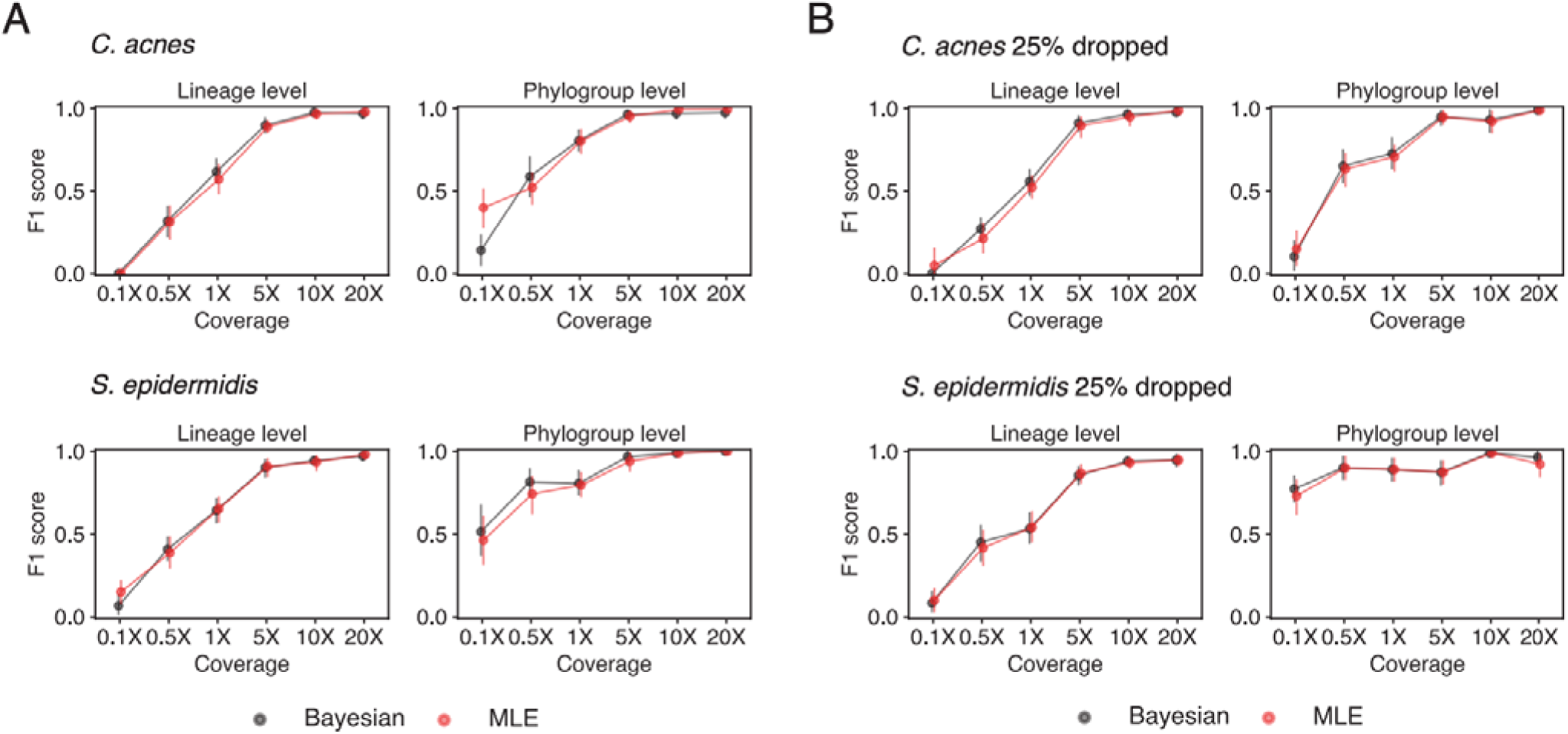
Minimal decrease in performance between Bayesian and Maximum Likelihood implementations of PHLAME model. We compared performance between Bayesian and maximum likelihood implementations of the PHLAME model (Supplemental Methods) using simulated metagenome benchmarks (see Fig. 3). In the Bayesian implementation, we estimated full posterior distributions over π and required 50% of the posterior distribution to be below 0.35 in order for a clade to count as detected. In the maximum likelihood implementation, we only inferred a point estimate on π and required this estimate to be below 0.35 in order for a clade to count as detected. Across species and simulations, results between the Bayesian and maximum likelihood implementations are largely consistent, with the maximum likelihood approach achieving only a minimal decrease in F1 score.

**Figure S6:**
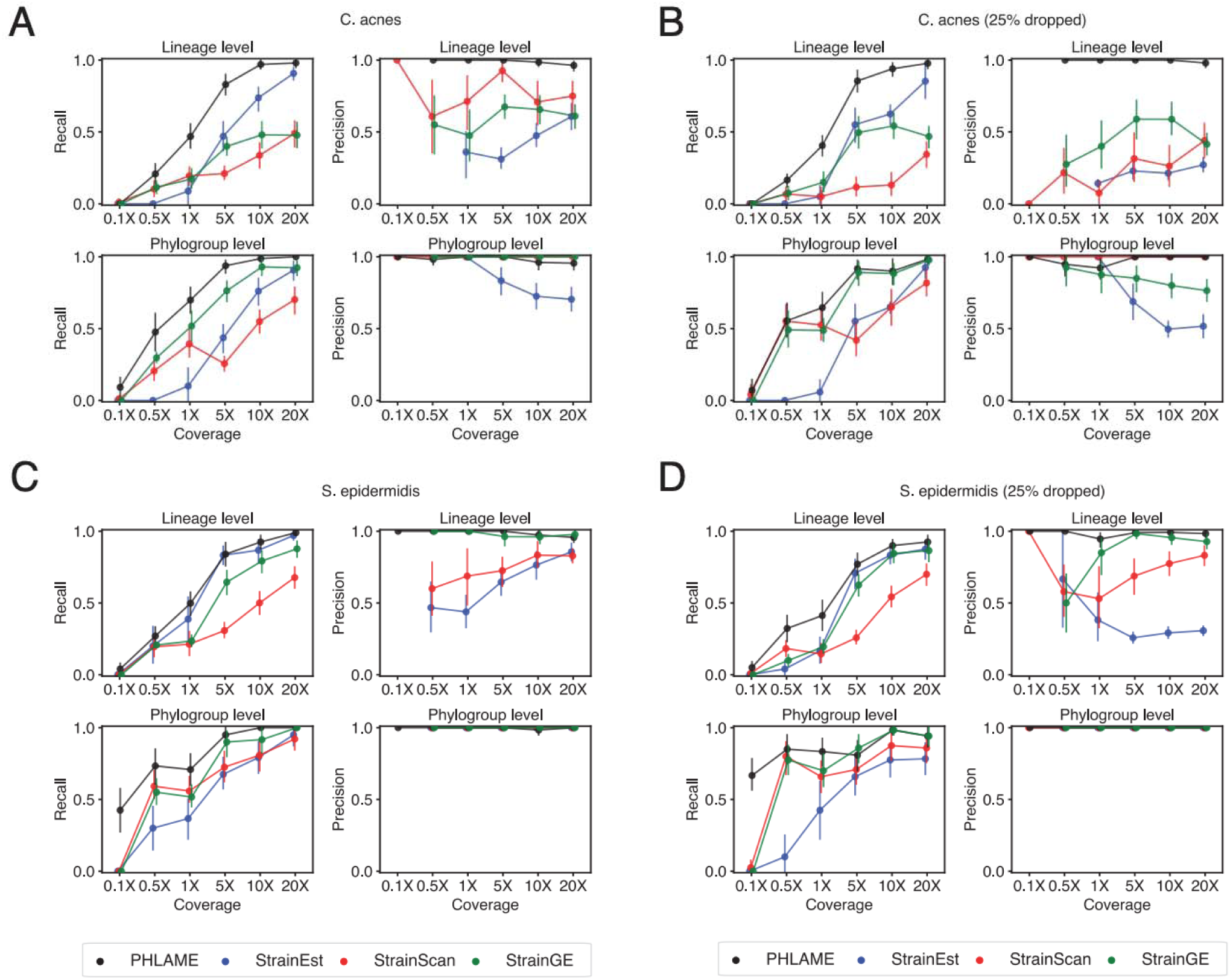
Precision and recall plots for synthetic metagenome benchmarks. Recall (left columns) and precision (right columns) for PHLAME, StrainEst, StrainGE, and StrainScan in synthetic metagenome benchmarks. Similar plots are shown for databases for (A) *C. acnes* with a perfect database; (B) *S. epidermidis* with a perfect database; (C) *C. acnes* with 25% of the clades held out; and (D) *S. epidermidis* with 25% of the clades held out. F1 scores are shown in Fig. 3. See also Fig. S7 and Fig. S8.

**Figure S7:**
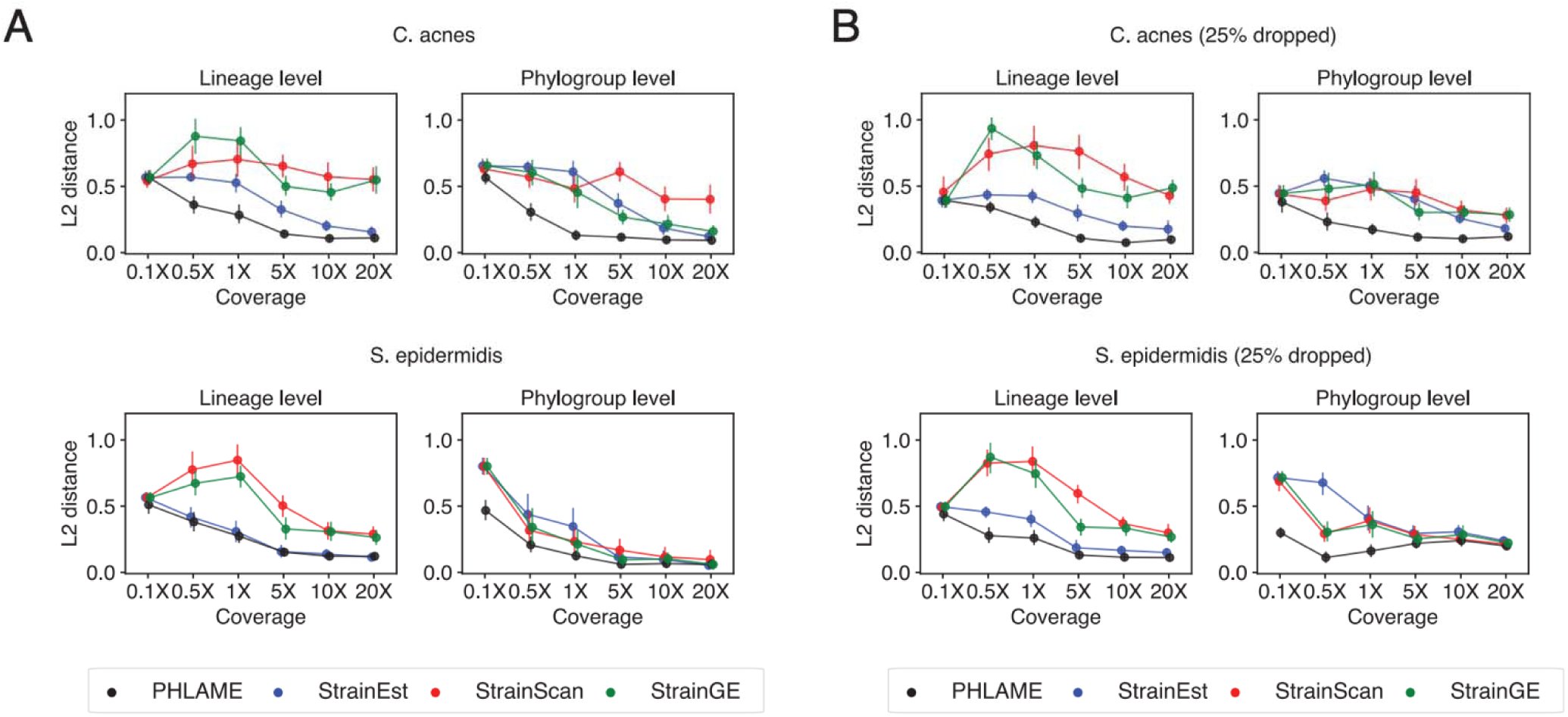
Relative abundance benchmarking using L2 distance. We measure accuracy in relative abundance estimates using L2 distance, which measures the Euclidean distance between the grou d truth relative abundance vector and a given estimated relative abundance vector (lower is better). (A) L2 distance calculated for different methods in simulated metagenome benchmarks with a perfect *C. acnes* or *S. epidermidis* reference database. (B) L2 distance calculated for different methods in simulated metagenome benchmarks where 25% of the phylogroups are randomly held out.

**Figure S8:**
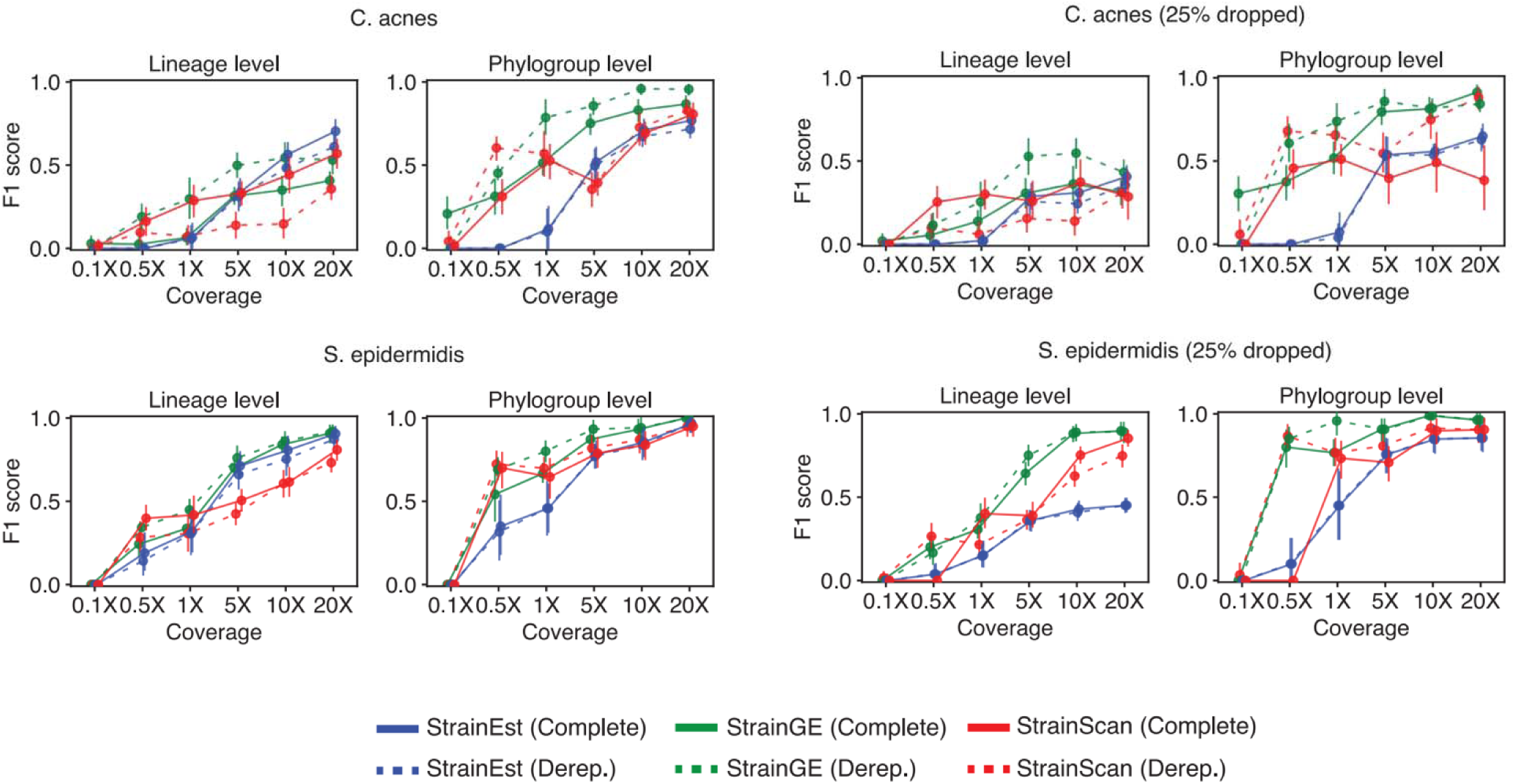
Performance between complete and dereplicated databases for StrainEst, StrainGE, and StrainScan. F1 score for each StrainEst, StrainGE, and StrainScan given either a complete database containing all reference genomes for a species (solid lines) or a dereplicated database containing a single representative genome per lineage (dashed lines).

**Figure S9:**
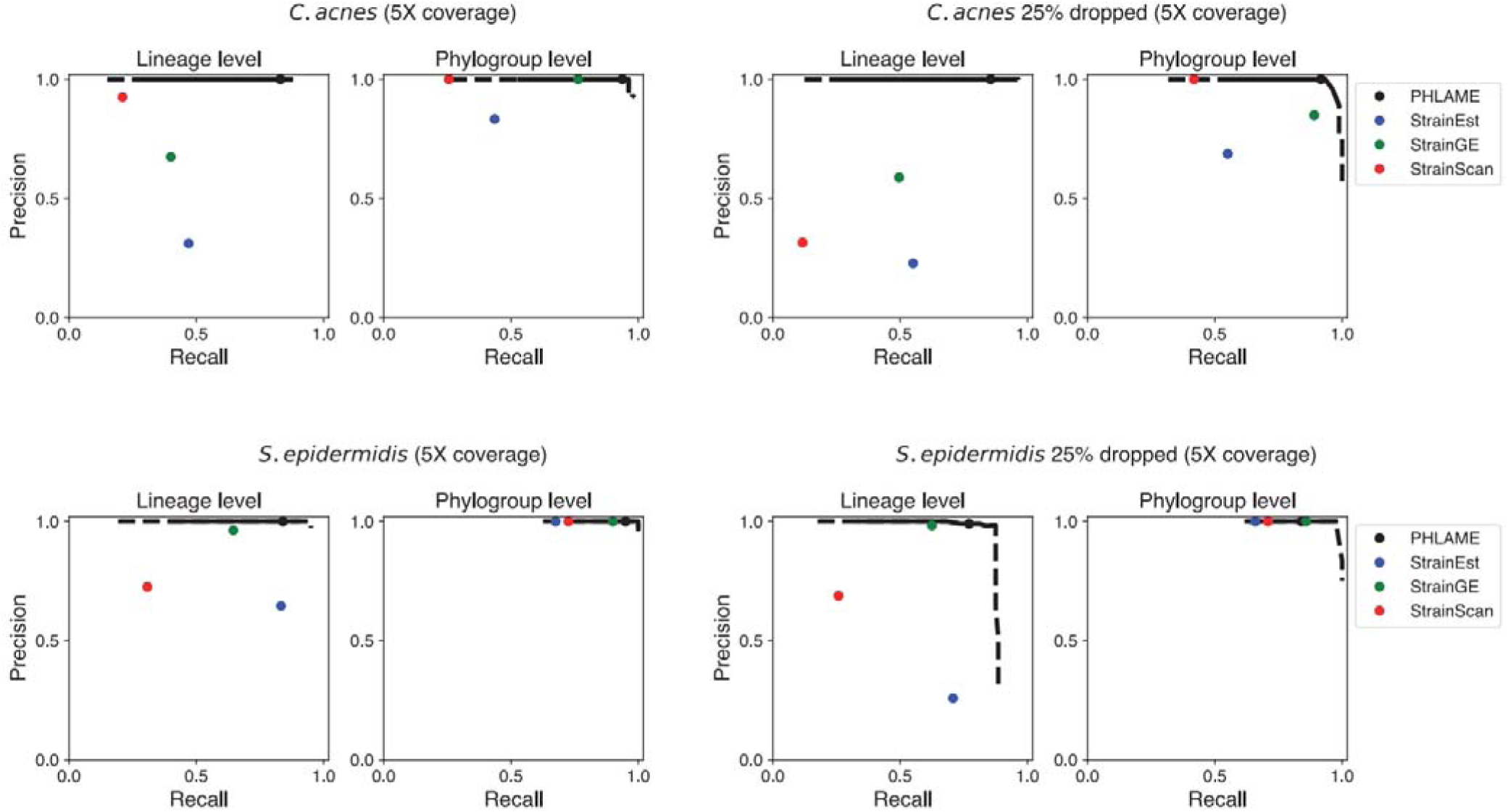
PHLAME results are robust to varying parameters. Precision-recall curves created by varying the detection threshold for PHLAME. The main detection threshold used for PHLAME requires at least 50% of the posterior density for π to be below a certain DV_b_ value. While possible values of DV_b_ range from 0 to 1, not all parameters are reasonable (for example, accepting detections when 50% of the posterior density for π is below 0.90 DV_b_ may be overly permissive). Focusing on the set of benchmarks where the focal species was subsampled to 5X coverage, we show ROC curves for PHLAME across the full range of possible π thresholds (0-1, dashed lines), as well as a set of possible π thresholds that we consider reasonable π thresholds in applied use (0.05-0.5, solid line). Mean precision and recall of StrainEst, StrainGE, and StrainScan for the same set of simulations are shown in colored dots, and mean precision and recall for the default PHLAME parameters shown in Figs. 3, S6-S7 is shown as a black dot.

**Figure S10:**
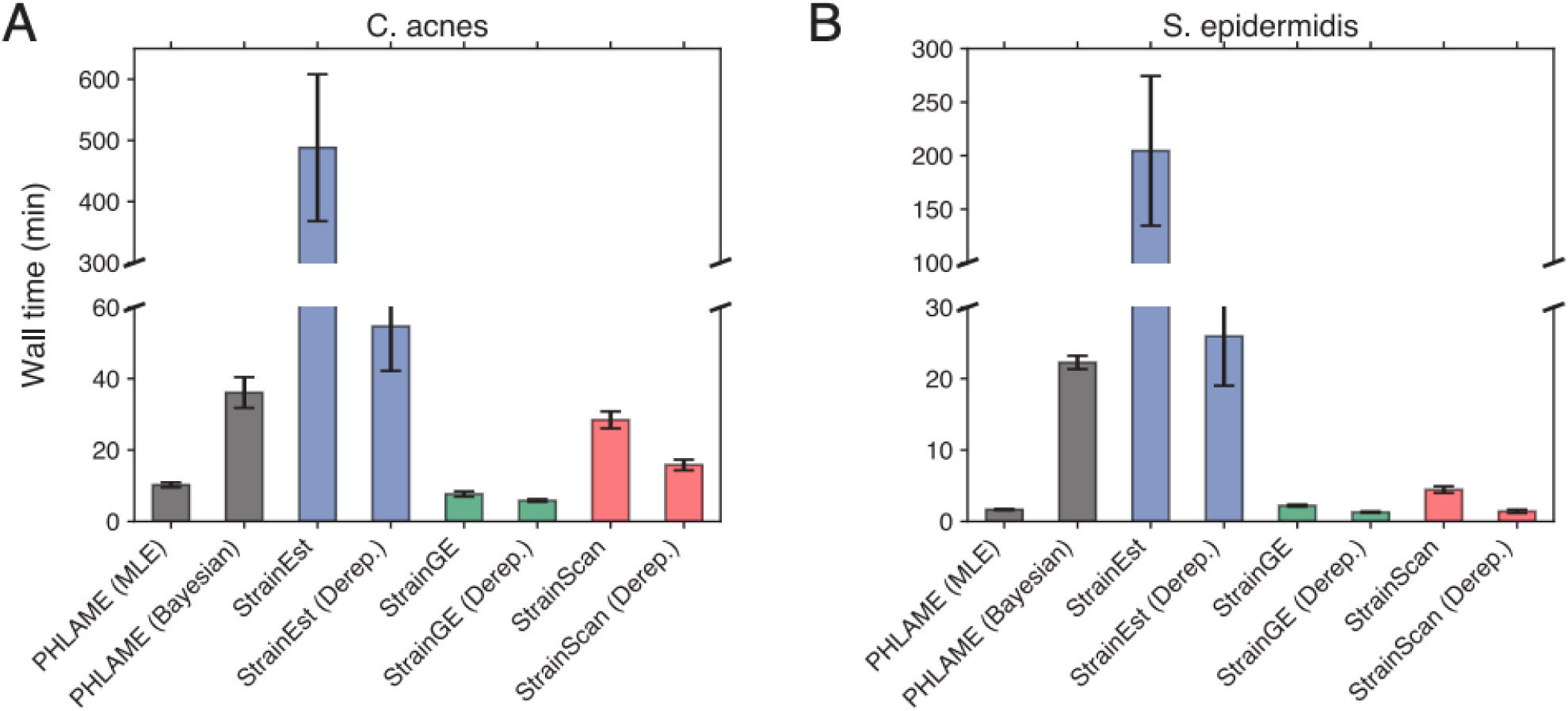
Comparison of method runtimes. Wall time (mean and 95% CI) to run PHLAME, StrainEst, StrainGE, and StrainScan. Read-to-result wall times were measured for (A) C. acnes and (B) S. epidermidis on the set of simulations where the focal species was at 20X coverage, and no genomes were held out from databases. For PHLAME and StrainEst, the time reported includes the time associated with read alignment and bam conversion. Each method was run on an AMD EPYC 7513 2.6GHz 64-core processor with 24Gb of allocated memory. Wall times were measured using the snakemake (v.7.20.0) benchmark utility.

**Figure S11:**
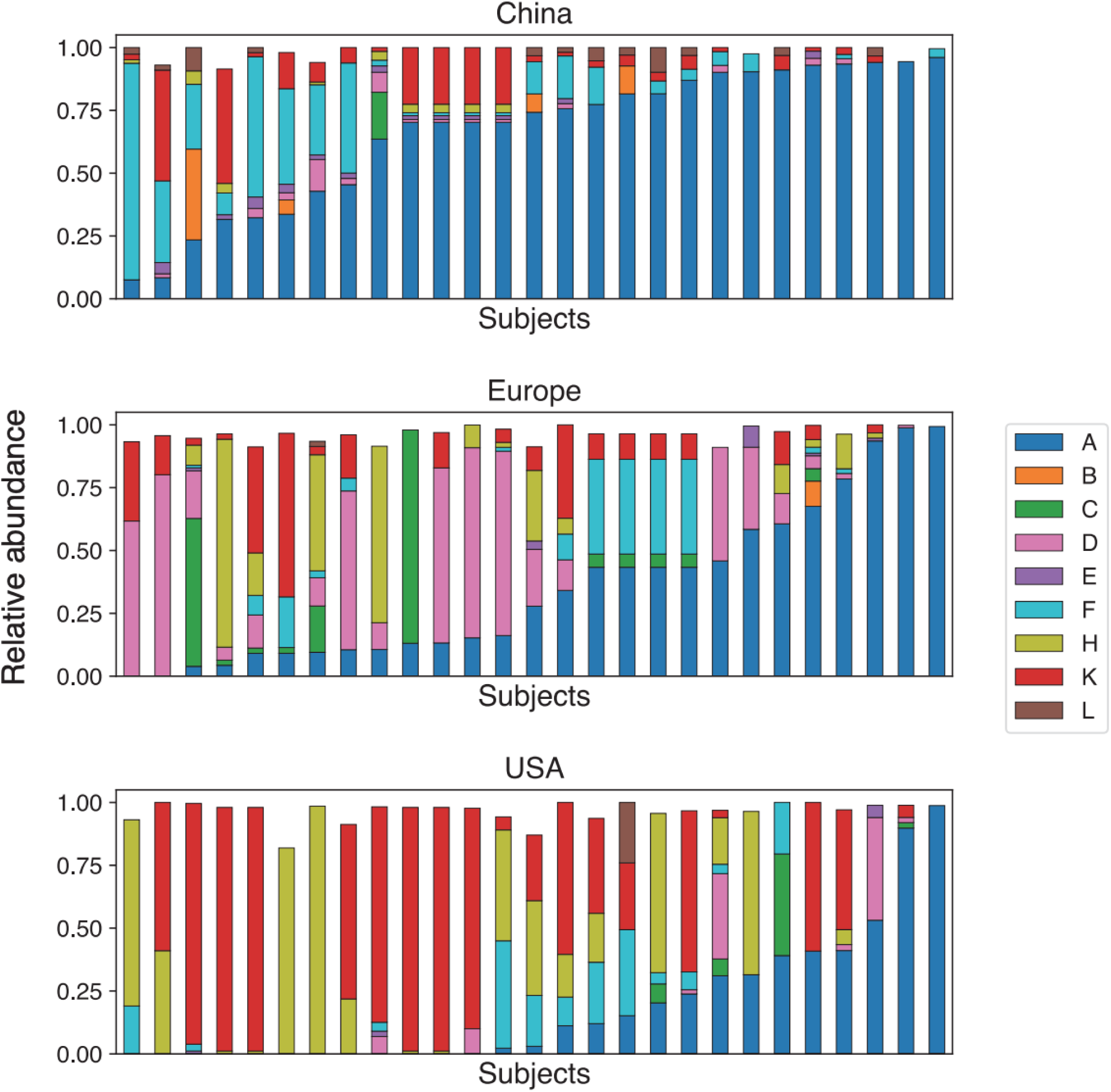
On-person taxonomic abundances of *C. acnes* phylogroups. 25 random taxonomic barplots per geographic region showing on-person *C. acnes* phylogroup abundances (one bar represents one subject).

**Figure S12:**
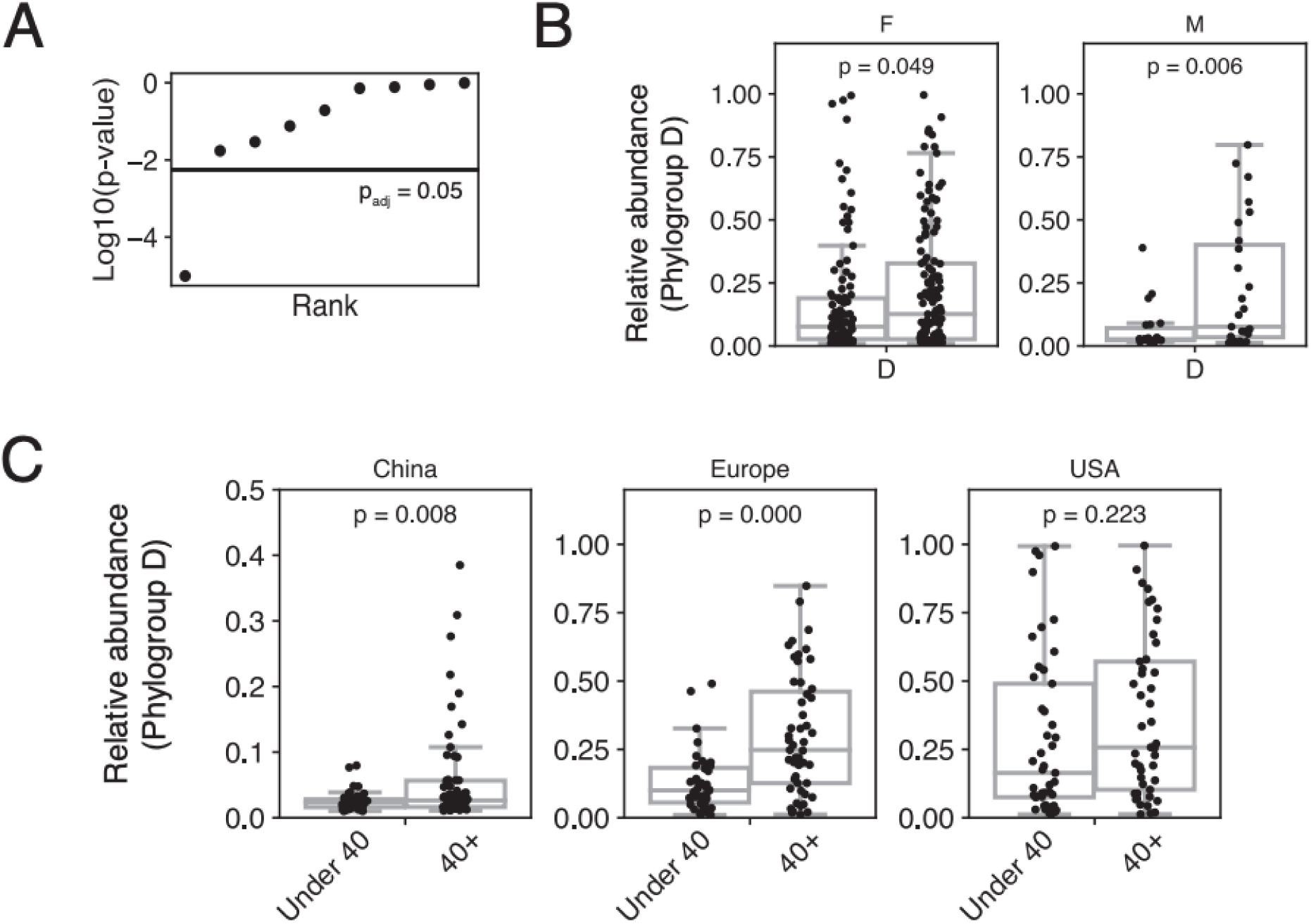
The relationship between *C. acnes* phylogroup D and age is robust across testable confounders. Multiple hypothesis correction for a rank-sum test for difference in phylogroup relative abundance between individuals under 40 compared to 40+. Black line represents an alpha of 0.05 after Bonferroni correction; only one phylogroup (phylogroup D) is significant after correction. (B) Difference in phylogroup D frequency on individuals under 40 compared to 40+, partitioned by reported sex. P-values represent the result of a Wilcoxon rank sum test. (C) Difference in phylogroup D frequency on individuals under 40 compared to 40+, partitioned by geographic region. P-values represent the result of a rank sum test. For the USA, this difference is not significant between individuals under 40 compared to 40+, but there is a significant rank correlation between phylogroup D and age (Fig. 5E).

**Figure S13:**
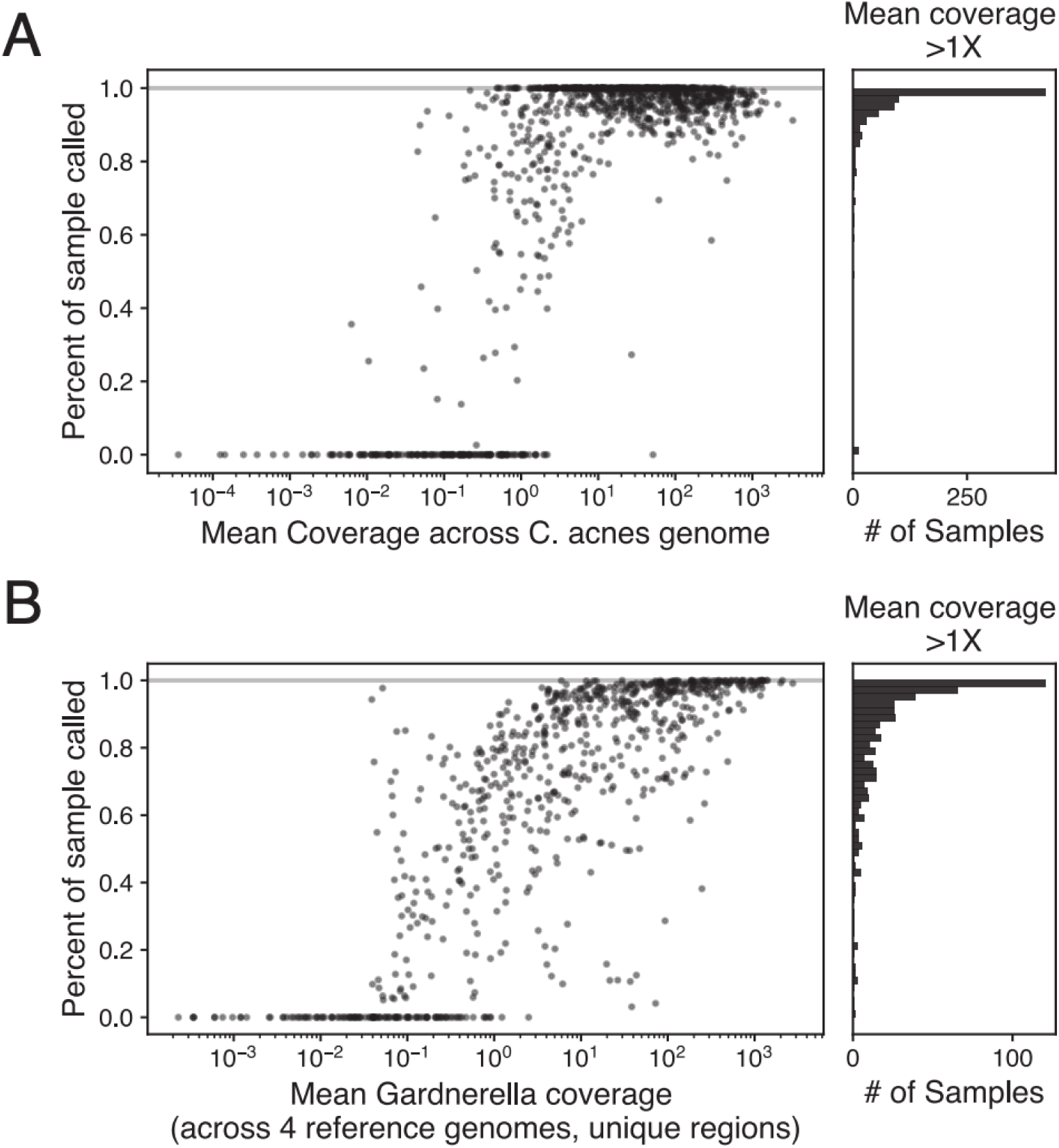
Novel *C. acnes* diversity is uncommon at the phylogroup level; novel *Gardnerella* diversity is more common. (A) Left: The percent of each sample classified by PHLAME at the phylogroup level as a function of mean coverage across the *C. acnes* genome (each dot represents one sample). Right: Histogram of the percent of sample classified by PHLAME at the phylogroup level, for only samples that reached greater than 1X mean coverage across the *C. acnes* genome. (B) Same diagram as (A) for *Gardnerella*. The coverage reported here is the mean coverage for each reference genome’s unique regions, summed across the four reference genomes used to classify *Gardnerella* diversity. A larger proportion of the Gardnerella diversity is unclassifiable in vaginal microbiome samples with greater than 1X coverage, compared to *C. acnes* diversity in skin microbiome samples (p < 0.001 two sample K-S test).

**Figure S14:**
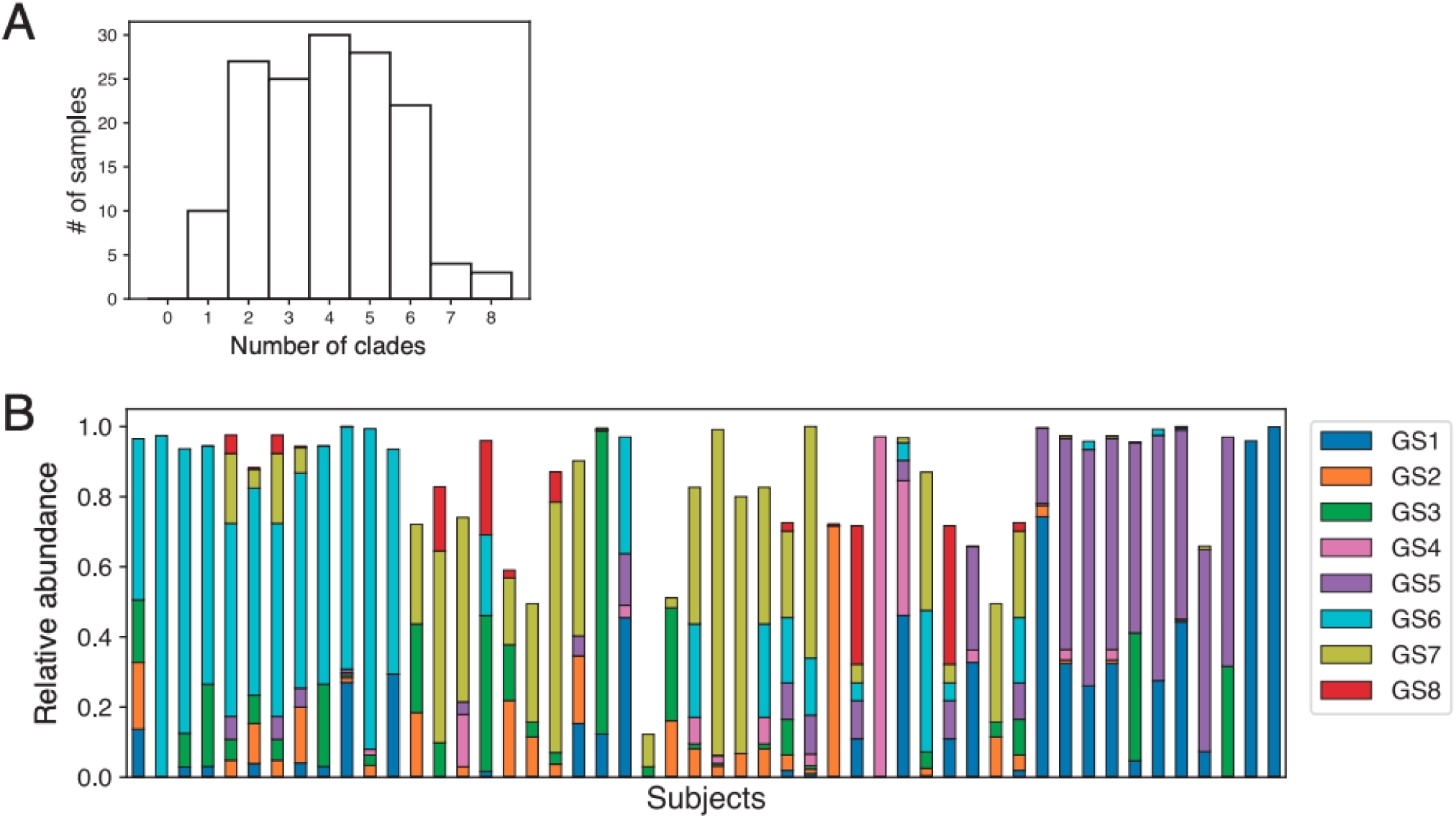
*Gardnerella* variation across subjects in the vaginal microbiome. (A) Number of *Gardnerella* clades (out of 8) per sample, detected across 149 subjects with > 3X coverage across 4 *Gardnerella* reference genomes (1 sample per subject). (B) Relative abundances of *Gardnerella* taxa in 50 random subjects with >3X coverage across 4 *Gardnerella* reference genomes (1 sample per subject).

**Figure S15:**
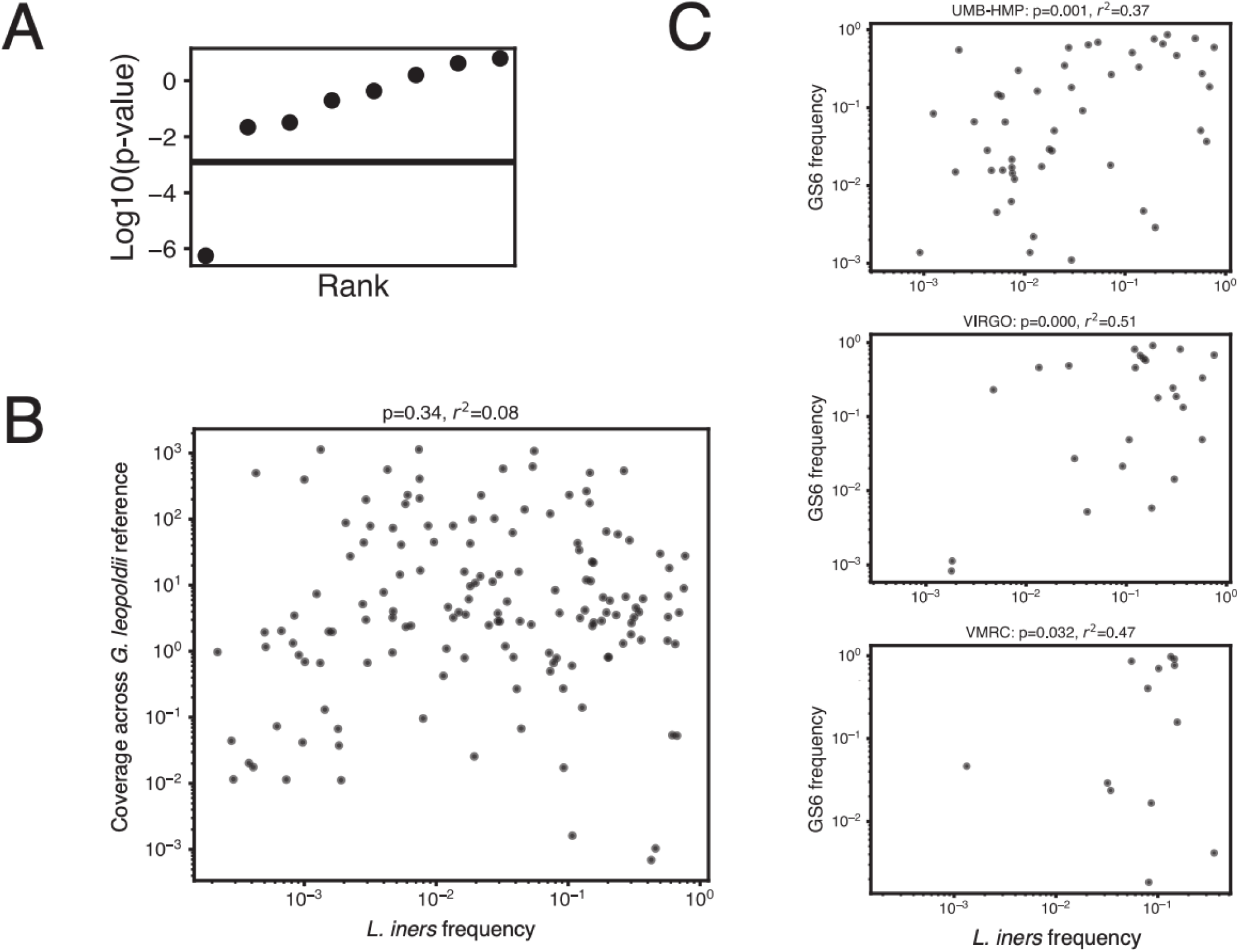
Association between GS6 (*G. swidsinskii*) and *L. iners* in the vaginal microbiome is consistent across possible confounders. (A) Multiple hypothesis correction (Spearman correlation) between *Gardnerella* clade frequency within the *Gardnerella* population and *L. iners* frequency in the sample. Black line represents a Bonferroni-corrected alpha of 0.01. After multiple hypothesis correction, only 1 clade (GS6) has a significant relationship with *L. iners* frequency. (B) There is no relationship between the frequency of *L. iners* in a sample and the number of reads mapping to the corresponding reference genome for GS6 (*G. leopoldii* 6420B), indicating that this association is not confounded by sequencing depth. Spearman’s correlation coefficient and significance value are shown above the plot. (C) Association between the relative frequency of GS6 in the *Gardnerella* population and *L. iners* frequency in the species-level community remains significant across different studies (UMB-HMP, VIRGO, VMRC).

**Figure S16:**
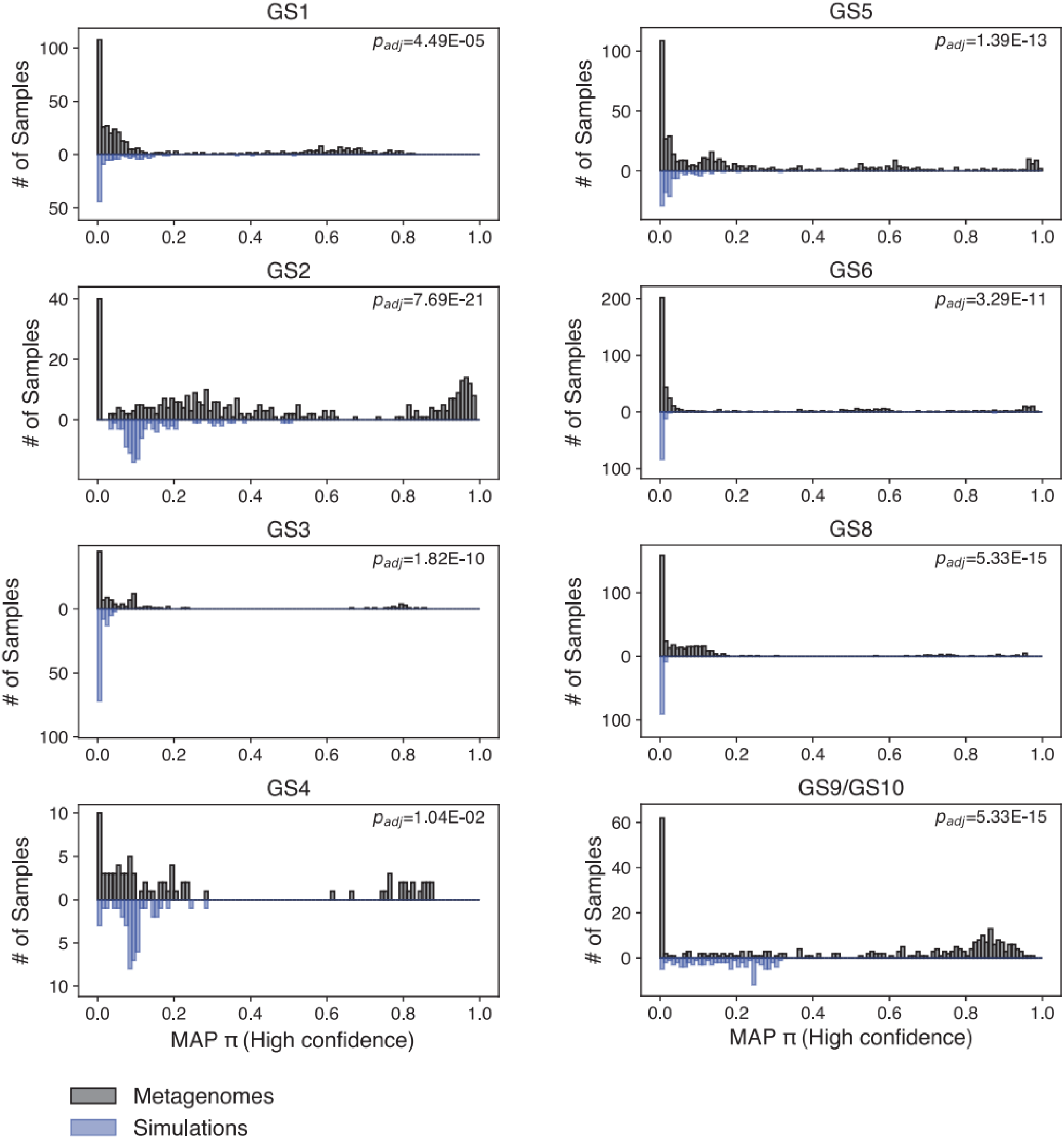
Peaks in high-confidence π values across samples suggest novel *Gardnerella* clades. Histograms showing the distribution of high-confidence maximum a posteriori (MAP) estimates for π, calculated with respect to each defined *Gardnerella* clade. MAP estimates for π are taken from real metagenomes (black) and 100 simulated metagenomes (blue) composed of random combinati ns of *Gardnerella* genomes. Results from a two-sample K-S test comparing the distribution of π estimates between real and simulated metagenomes are shown next to each graph (Bonferroni-corrected p-values).

**Figure S17:**
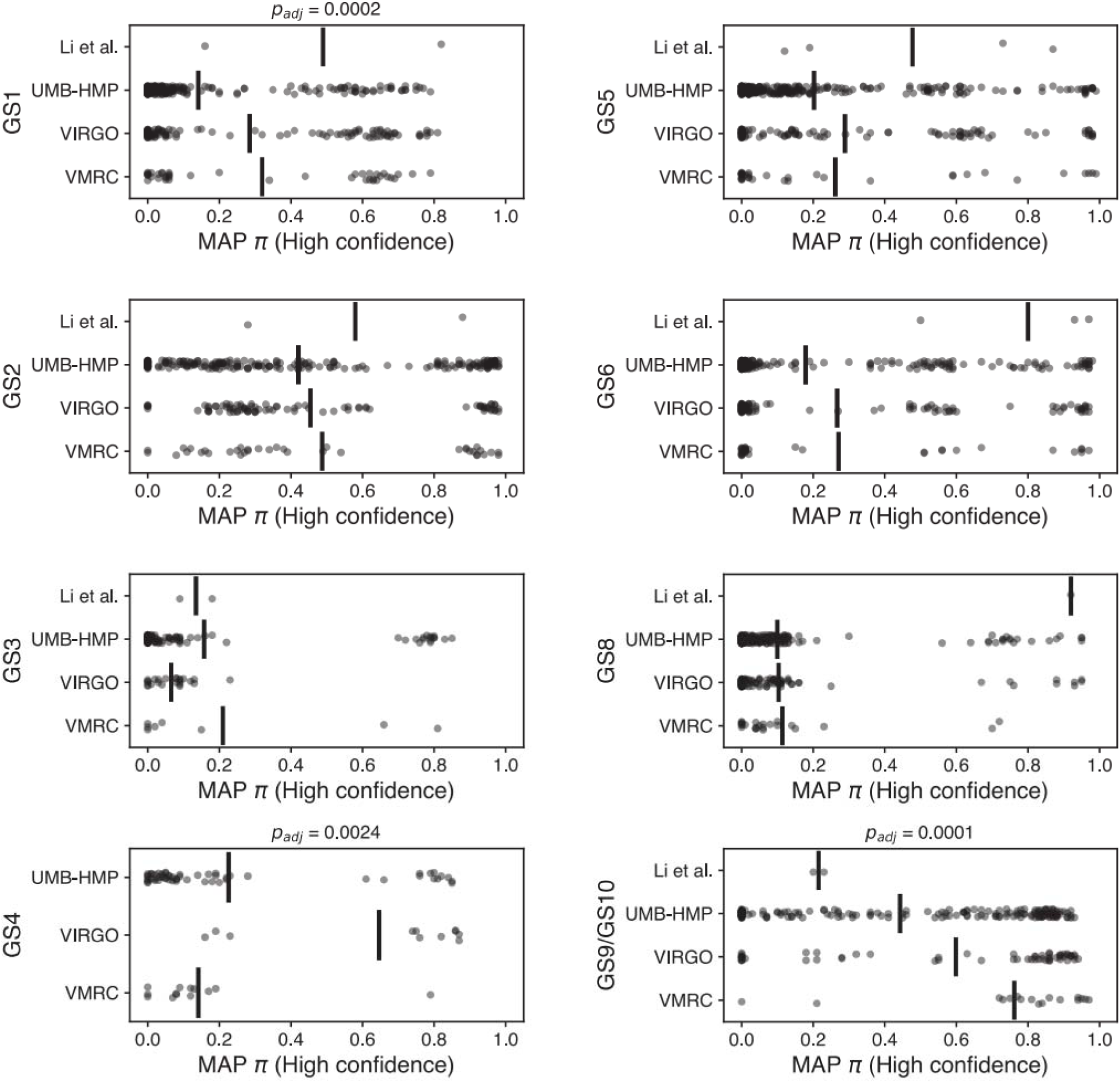
Putative novel *Gardnerella* clades are enriched in specific studies and samples. MAP estimates for π, calculated with respect to each defined *Gardnerella* clade, separated by study. Each dot represents one estimate from one sample; mean lines for studies are shown in bars. Adjusted p-values for clades that have significantly differently distributed π estimates across studies (Kruskal-Wallis test, Bonferroni-corrected).

