## Supplemental Text for "Intraspecies associations from strain-rich metagenome samples"

#### 1 Posterior Inference Algorithm

We implement a Slice-within-Gibbs sampler which offers quick runtime and removes the need to select free sampling parameters. PHLAME models the number of reads supporting a clade-specific allele ($x_i$) at informative position $i$ as a random variable originating from a zero-inflated negative binomial distribution. We can describe this distribution with the following hierarchical model:

$$x_i \sim Poisson(\lambda_i z_i)$$

$$z_i \sim Bernoulli(1-\pi)$$

$$\lambda_i \sim Gamma(\alpha, \beta)$$

We would like to recover the posterior distribution $f(\pi, \alpha, \beta | x_i)$, which will give us information about both the divergence ($DV_b$) in a sample (equivalent to $\pi$), as well as the expected value of our counts (equivalent to $\alpha/\beta$). We can start with the likelihood function for observed counts $x_i$, which is simply a Poisson likelihood:

$$f(x_i|\lambda_i z_i) = \prod_{i=1}^{n} \frac{(\lambda_i z_i)^{x_i} e^{-\lambda_i z_i}}{x_i!} \quad (1)$$

Using Bayes rule, we can write the full posterior for $f(x_i|\lambda_i, z_i)$ as a product of the likelihood and the prior probability:

$$f(\lambda_i, z_i|x_i) = \frac{f(x_i|\lambda_i, z_i) \times f(\lambda_i, z_i|\pi, \alpha, \beta)}{f(x_i)} \quad (2)$$

The distribution of the prior probability $f(\lambda_i, z_i|\pi, \alpha, \beta)$ is conditional on several variables we have defined in our model. We can thus write the full form of the prior as follows:

$$f(\lambda_i, z_i|\pi, \alpha, \beta) = f(\lambda_i|\alpha, \beta) f(z_i|\pi) f(\alpha, \beta|m, v, s, p) f(\pi) \quad (3)$$

For the Gamma distribution $\lambda_i \sim Gamma(\alpha, \beta)$, we use a conjugate prior of the form $f(\alpha, \beta) = \frac{\beta^{v\alpha}}{\Gamma(\alpha)^m} p^{\alpha-1} e^{-s\beta}$ with four free parameters: $m, v, s, p$ [1]. This will allows us recover most conditional distributions as standard probability distributions. For the prior distribution $f(\pi)$ we use $Uniform[0, 1]$ for simplicity, although this in theory could take a different form to encode prior information on the distribution of $\pi$. We can rewrite each conditional probability as the analytical form of the probability distribution described in our model.

$$f(\lambda_i, z_i) = \left( \prod_{i-1}^{n} \frac{1}{\Gamma(\alpha)} \lambda_i^{\alpha-1} e^{-\beta\lambda_i} \times \pi^{z_i} 1 - \pi^{1-z_i} \right) \times \frac{\beta^{v\alpha}}{\Gamma(\alpha)^m} p^{\alpha-1} e^{-s\beta} \times 1$$

$$\propto \frac{\beta^{\alpha(n+v)}}{\Gamma(\alpha)^{m+n}} e^{-\beta(\sum^{n} \lambda_i)} \pi^{nz_i} (1-\pi)^{n-nz_i} p^{\alpha-1} \prod_{i=1}^{n} \lambda_i^{\alpha-1} \quad (4)$$

The full posterior distribution is proportional to the product of (1) and (4):

$$f(\lambda_i, z_i|x_i) \propto \frac{\beta^{\alpha(n+v)}}{\Gamma(\alpha)^{m+n}} e^{-\beta(\sum^n \lambda_i)-\sum^n \lambda_i z_i} \pi^{\sum^n z_i}(1-\pi)^{n-\sum^n z_i} p^{\alpha-1} \prod_{i=1}^n \lambda_i^{a-1}(\lambda_i z_i)^{x_i} \tag{5}$$

From (5), we can derive the conditional distribution for each parameter by taking only the terms dependent on the parameter.

$$\begin{aligned}
f(\lambda_i|...) &\propto e^{-\beta(s+\sum^n \lambda_i)-\sum^n \lambda_i z_i} \prod_{i=1}^n \lambda_i^{\alpha-1}(\lambda_i z_i)^{x_i} \\
&\propto e^{-\beta(s+\lambda_i)-\sum^n \lambda_i z_i} \lambda_i^{x_i+\alpha-1} \prod_{i=1}^n z_i^{y_i} \\
&\propto \begin{cases} z_i = 0; & e^{-\beta(s+\lambda_i)} \lambda_i^{x_i+\alpha-1} \prod_{i=1}^n 0^{y_i} \\ z_i = 1; & e^{-\beta(s+\lambda_i)-\sum^n \lambda_i} \lambda_i^{x_i+\alpha-1} \prod_{i=1}^n 1^{y_i} \end{cases}
\end{aligned}$$

$$\begin{aligned}
f(\lambda_i|z_i = 0, ...) &\propto \begin{cases} x_i = 0; & e^{-\beta(s+\lambda_i)} \lambda_i^{x_i+\alpha-1} \\ x_i > 0; & 0 \end{cases} \\
&\propto Gamma(\alpha + x_i, \beta); s = 0
\end{aligned}$$

$$\begin{aligned}
f(\lambda_i|z_i = 1, ...) &\propto \begin{cases} x_i = 0; & e^{-\beta(s+\lambda_i)-\lambda_i} \lambda_i^{\alpha-1} \\ x_i > 0; & e^{-\beta(s+\lambda_i)-\lambda_i} \lambda_i^{x_i+\alpha-1} \end{cases} \\
&\propto Gamma(\alpha + x_i, \beta + 1); s = 0
\end{aligned} \tag{6}$$

$$\begin{aligned}
f(z_i|...) &\propto e^{-\sum^n \lambda_i} \pi^{-\sum^n z_i}(1-\pi)^{n-\sum^n z_i} \prod_{i=1}^n z_i^{x_i} \\
&\propto \frac{e^{-\sum^n \lambda_i} \pi^{-\sum^n z_i}}{(1-\pi)^{z_i}} z_i^{x_i} \\
f(z_i|x_i = 0, ...) &\propto \frac{e^{-\sum^n \lambda_i z_i} \pi^{z_i}}{(1-\pi)^{z_i}} \\
&\propto \begin{cases} z_i = 1; & \frac{e^{-\sum^n \lambda_i} \pi}{1-\pi} = \frac{e^{-n\bar{\lambda}\pi}}{e^{-n\bar{\lambda}}\pi+(1-\pi)} \\ z_i = 0; & 1 - \frac{e^{-n\bar{\lambda}\pi}}{e^{-n\bar{\lambda}}\pi+(1-\pi)} = \frac{1-\pi}{e^{-n\bar{\lambda}}\pi+(1-\pi)} \end{cases} \\
f(z_i|x_i > 0, ...) &\propto \begin{cases} z_i = 1; & \frac{e^{-n\bar{\lambda}}\pi}{e^{-n\bar{\lambda}}\pi+(1-\pi)I(x_i=0)} \\ z_i = 0; & 1 - \frac{e^{-n\bar{\lambda}}\pi}{e^{-n\bar{\lambda}}\pi+(1-\pi)I(x_i=0)} = \frac{e^{-n\bar{\lambda}}\pi+(1-\pi)I(x_i=0)-e^{-n\bar{\lambda}}\pi}{e^{-n\bar{\lambda}}\pi+(1-\pi)I(x_i=0)} \end{cases} \\
&\propto Bernoulli\left(\frac{e^{-n\bar{\lambda}}\pi}{e^{-n\bar{\lambda}}\pi + (1-\pi)I(x_i = 0)}\right)
\end{aligned} \tag{7}$$

$$
\begin{aligned}
f(\pi|...) &\propto \pi^{\sum^{n} z_i}(1-\pi)^{n-\sum^{n} z_i} \\
&\propto Beta\left(\sum^{n} z_i + 1, (n - \sum^{n} z_i) + 1\right)
\end{aligned} \tag{8}
$$

$$
\begin{aligned}
f(\alpha|...) &\propto \frac{\beta^{\alpha(n+v)}}{\Gamma(\alpha)^{m+n}} p^{\alpha-1} \prod_{i=1}^{n} \lambda_i^{a-1} \\
&\propto \frac{\beta^{\alpha(n+v)}}{\Gamma(\alpha)^{m+n}} \left(p \prod_{i=1}^{n} \lambda_i\right)^{\alpha} \\
&\Rightarrow a(n+v)ln(\beta) + \alpha\left(ln(p) + \sum^{n} ln(\lambda_i)\right) - (m+n)ln(\Gamma(\alpha))
\end{aligned} \tag{9}
$$

$$
\begin{aligned}
f(\beta|...) &\propto \beta^{\alpha(n+v)-1} e^{-\beta(s+\sum^{n}\lambda_i)} \\
&\propto Gamma\left(\alpha(n+v), s + \sum^{n} \lambda_i\right)
\end{aligned} \tag{10}
$$

Thus, all but one conditional can be recovered as standard, easy to sample distributions:

- $f(\lambda_i|...)$ can be sampled as $Gamma(\alpha + x_i, \beta + z_i)$.
- $f(z_i|...)$ can be sampled as $Bernoulli(\frac{e^{-n\bar{\lambda}}\pi}{e^{-n\bar{\lambda}}\pi + (1-\pi)I(x_i=0)})$, where $I(x_i = 0)$ is the indicator function.
- $f(\pi|...)$ can be sampled as $Beta\left(\sum^{n} z_i + 1, (n - \sum^{n} z_i) + 1\right)$
- $f(\beta|...)$ can be sampled as $Gamma\left(\alpha(n+v), s + \sum^{n} \lambda_i\right)$
- $f(\alpha|...)$ does not take the form of a standard distribution; we can instead use a Slice sampler to take from the ln conditional $a(n+v)ln(\beta) + \alpha\left(lnp + \sum^{n} ln(\lambda_i)\right) - (m+n)ln(\Gamma(\alpha))$.

Information on the overall dispersion of reads covering a set of positions can be encoded in the prior distribution over $Gamma(\alpha, \beta)$. First, we take $y_i$ to be the number of reads supporting alleles in informative positions $i$. We assume that $y_i$ originates from an ordinary negative binomial distribution, which we will consider in the parameterization $y_i \sim NB(\mu, \alpha)$, where $\mu$ is the rate parameter and $\alpha$ is the dispersion relative to a Poisson and equivalent to the first parameter from the $Gamma(\alpha, \beta)$ distribution. The maximum likelihood estimates of both parameters have an analytical solution:

$$
\begin{aligned}
\hat{\mu} &= \frac{\sum^{n} y_i}{n} \\
\hat{\alpha} &= \frac{\hat{\mu}^2}{Var(y_i) - \hat{\mu}}
\end{aligned} \tag{11}
$$

If we observe the form of our conjugate prior, it becomes apparent that if we take $s = 0, v = 0$, the impact of the prior on $\beta$ is negligible. We can therefore simplify our prior distribution.

$$\begin{aligned} f(\alpha, \beta) &= \frac{\beta^{v\alpha}}{\Gamma(\alpha)^m} p^{\alpha-1} e^{-s\beta} \\ &\rightarrow \frac{1}{\Gamma(\alpha)^m} p^{\alpha-1} \end{aligned} \tag{12}$$

In this form, $p$ informs the mode of the distribution while $m$ controls the relative strength of the prior. PHLAME takes $m = 20$ by default. We can create priors centered around certain values of $\alpha$ as follows,

$$\begin{aligned} m &= 20 \\ ln(p) &= m \times \psi(\hat{\alpha}) \end{aligned} \tag{13}$$

where $\psi$ is the digamma function. Note that we need only specify $ln(p)$ because we are sampling from the log conditional of $f(\alpha|...)$.

### 2 Maximum Likelihood Implementation

The maximum likelihood implementation of the PHLAME model is simpler, and requires minimizing the log-likelihood function while $\alpha = \hat{\alpha}$. The log-likelihood function for a zero-inflated negative binomial distribution with parameters $\pi$ (zero inflation), $\mu$ (rate), and $\alpha$ (overdispersion) is:

$$\ell(\pi, \mu, \alpha | x_i) = \sum_{i=1}^{n} \begin{cases} ln\left(\pi + (1-\pi)\left(1 - \frac{\alpha}{\mu+\alpha}\right)^{\alpha}\right) & x_i = 0 \\ \sum_{i=1}^{n} ln(1-\pi) + ln\left(\frac{\Gamma(x_i+\alpha)}{x_i!\Gamma(\alpha)}\right) + \alpha ln\left(\frac{\mu}{\mu+\alpha}\right) + x_i ln\left(\frac{\alpha}{\mu+\alpha}\right) & x_i > 0 \end{cases}$$

We can then minimize over $\pi$ and $\mu$, while keeping $\alpha$ constant at $\hat{\alpha}$:

$$\hat{\pi}, \hat{\mu} = \arg\min_{\pi,\mu} -\ell(\pi, \mu, \hat{\alpha} \mid y_1, \ldots, y_n)$$
